# *Txikispora philomaios* n. sp., n. g., a Micro-Eukaryotic Pathogen of Amphipods, Reveals Parasitism and Hidden Diversity in Class Filasterea

**DOI:** 10.1101/2021.01.19.427289

**Authors:** Ander Urrutia, Konstantina Mitsi, Rachel Foster, Stuart Ross, Martin Carr, Ionan Marigomez, Michelle M. Leger, Iñaki Ruiz-Trillo, Stephen W. Feist, David Bass

## Abstract

This study provides a morphological, ultrastructural, and phylogenetic characterization of a novel micro-eukaryotic parasite (2.3-2.6 µm) infecting genera *Echinogammarus* and *Orchestia*. Longitudinal studies across two years revealed that infection prevalence peaked in late April and May, reaching 64% in *Echinogammarus* sp. and 15% in *Orchestia* sp., but was seldom detected during the rest of the year. The parasite infected predominantly haemolymph, connective tissue, tegument, and gonad, although hepatopancreas and nervous tissue were affected in heavier infections, eliciting melanization and granuloma formation. Cell division occurred inside walled parasitic cysts, often within host haemocytes, resulting in haemolymph congestion. Small subunit (18S) rRNA gene phylogenies including related environmental sequences placed the novel parasite as a highly divergent lineage within Class Filasterea, which together with Choanoflagellatea represent the closest protistan relatives of Metazoa. We describe the new parasite as *Txikispora philomaios* n. sp. n. g., the first confirmed parasitic filasterean lineage, which otherwise comprises four free-living flagellates and a rarely observed endosymbiont of snails. Lineage-specific PCR probing of other hosts and surrounding environments only detected *T. philomaios* in the platyhelminth *Procerodes* sp. We expand the known diversity of Filasterea by targeted searches of metagenomic datasets, resulting in 13 previously unknown lineages from environmental samples.

## INTRODUCTION

The Class Filasterea Cavalier-Smith 2008 currently comprises five species (Shalchian-Tabrizi et al. 2008; Hehenberger et al. 2017; Tikhonenkov et al. 2020). Initially classified as a nucleariid, *Capsaspora owczarzaki* was the first filasterean to be described (Stibbs et al. 1979; Owczarzak et al. 1980; Amaral-Zettler et al. 2001; Hertel et al. 2002; Ruiz-Trillo et al. 2004). This filopodial amoeba is a facultative endosymbiont (Harcet et al. 2016) isolated from explanted pericardial sacs of laboratory-grown *Biomphalaria* sp. snails (Stibbs et al. 1979; Morgan et al. 2002), which remains elusive in environmental samplings (Hertel et al. 2004; del Campo and Ruiz-Trillo 2013; Shanan et al. 2015; Ferrer-Bonet and Ruiz-Trillo 2017; Arroyo et al. 2018). In contrast, the other four species (*Ministeria vibran*s, *Ministeria marisola, Pigoraptor chileana*, and *Pigoraptor vietnamica*) are free-living flagellates, sampled from marine and freshwater ecosystems (Patterson et al. 1993; Tong et al. 1997; Hehenberger et al. 2017; Mylnikov et al. 2019). The discovery of *C. owczarzaki* drew considerable scientific attention, as resistant cysts present in the mantle of *Biomphalaria glabrata* were observed to attack and kill sporocysts of the trematode *Schistosoma mansoni* parasitizing the snail (Stibbs et al. 1979; Eveland and Haseeb 2011). *S. mansoni*, which has *B. glabrata* as intermediate host, causes schistosomiasis in humans, a disease affecting over 230 million people worldwide (Colley et al. 2014).

Filasterea are also of interest (Ruiz-Trillo et al. 2008; Suga et al. 2013; Torruella et al. 2015; Hehenberger et al. 2017), as they branch phylogenetically close to the metazoan radiation, being sister to Choanozoa (the Metazoa + Choanoflagellatea clade) (Shalchian-Tabrizi et al. 2008; Paps et al. 2013; Torruella et al. 2015; López-Escardó et al. 2019). Morphological (James-Clark 1868), ultrastructural (Laval 1971; Hibberd 1975), and phylogenetic inference (Cavalier-Smith 1993; Wainright et al. 1993; Snell et al. 2001; King 2004; Ruiz-Trillo et al. 2006) suggested a common evolutionary origin for Metazoa and Choanoflagellatea, which was confirmed by phylogenomic analyses (King et al. 2005; Steenkamp et al. 2006; Ruiz-Trillo et al. 2008). Phylogenomic studies also revealed the relationship between genera *Capsaspora* and *Ministeria* and their sister-clade relationship to Choanozoa (Shalchian-Tabrizi et al. 2008 Torruella et al. 2012; Hehenberger et al. 2017). Since then, the genomes and transcriptomes of filasterean species have been thoroughly investigated to comprehend the evolutionary processes that drove the inception of animal multicellularity (Suga et al. 2013; Torruella et al. 2015; Sebé-Pedrós et al. 2017; Hehenberger et al. 2017; Grau-Bove et al. 2017).

For almost 40 years, our knowledge of filasterean ultrastructure came from a single paper (Owczarzak et al. 1980), describing *C. owczarzaki*. Recently, the ultrastructures of *M. vibrans* and *Pigoraptor* sp. have been investigated (Torruella et al. 2015; Mylnikov et al. 2019; Tikhonenkov et al. 2020). Regarding the ecology and global distribution of the species within the Class, existing information is limited to the sampling locations of type species, and some feeding observations under culture conditions (Stibbs et al. 1979; Tong 1997; Hehenberger et al. 2017; Mylnikov et al. 2019; Tikhonenkov et al. 2020). Given the low number of species described the influence of filastereans in the food web has been thought to be insignificant, at least in comparison to much bigger protistan clades, or notorious pathogenic taxa. Recent environmental studies have suggested the relationship between an abundant clade of marine opisthokonts (MAOP-1) and Filasterea (del Campo et al. 2015; Hehenberger et al. 2017; Heger et al. 2018), challenging the idea of a small and scarce group. Excluding the facultative endosymbiont *C. owczarzaki*, all filastereans and choanoflagellates are free-living organisms, contrasting with the parasitic lifestyle of ichthyosporeans (mesomycetozoeans) (Mendoza et al. 2002; Glockling et al. 2013), which includes important pathogens of fish (Ragan et al. 1996; Pekkarinen and Lotman 2003; Andreou et al. 2011), amphibians (Broz and Privora 1952; Pereira et al. 2005; Rowley et al. 2013), birds, and mammals, including humans (Fredricks et al. 2000; Silva et al. 2005).

During a histopathological survey of invertebrates inhabiting the intertidal zone (Weymouth, UK), an unidentified protist was observed parasitizing two of the most common species of amphipods (*Echinogammarus* sp. and *Orchestia* sp.). Analysis by light microscopy of the structure and tissue tropism of the parasite did not allow a clear assignment of the organism to any of the pathogen groups commonly observed infecting amphipods or crustaceans. Similarly, examination of the ultrastructure by transmission electron microscopy (TEM), did not show any distinctive organelle suggesting taxonomic affiliation. Preliminary phylogenetic analyses of the 18S SSU rRNA strongly indicated that this lineage was a highly divergent novel genus within Holozoa. However, it did not consistently branch with any of the four established unicellular clades (Choanoflagellatea, Filasterea, Corallochytrea/Pluriformea, and Ichthyosporea/ Mesomycetozoea). When a greater diversity of environmental holozoan sequences were included in the analyses the parasite branched with Filasterea as the earliest diverging branch. This study comprises a complete histopathological, ultrastructural, and phylogenetic analysis based on the complete 18S SSU rRNA of the novel parasite, described as *Txikispora philomaios* n. sp. n. g. We also present data on its prevalence, host range, biological cycle, and potential transmission routes. Additionally, we demonstrate novel filasterean diversity on the basis of sequences mined from environmental sequencing datasets. The description of *T. philomaios* and its parasitic lifestyle adds to a growing understanding of filasterean diversity, ecology, and lifestyle traits.

## MATERIALS AND METHODS

### Sample collection

Amphipods belonging to genera *Orchestia, Echinogammarus, Gammarus*, and *Melita* were collected in the Tamar estuary (Torpoint, Cornwall), Camel estuary (Padstow, Cornwall), Dart estuary (Dittisham, Devon) and Newton’s Cove (Weymouth, Dorset, UK) between 2016 and 2018 (Table 1; Fig. 1). Individuals *of Echinogammarus* sp. and *Orchestia* sp. were sampled in the upper part of the intertidal zone, behind rocks and algae. Individuals of *Gammarus* sp. and *Melita* sp. were sampled in the lower part of the intertidal behind small stones and submerged algae. In addition to amphipods, other very abundant invertebrates sharing the same habitat in the upper part of the intertidal were also collected in Newton’s Cove from May 2019 to September 2019 (Table 2). These organisms include *Capitella* sp. (Polychaeta, Annelida), *Procerodes* sp. (Turbellaria, Platyhelminthes) and harpacticoid copepods of the Ameiridae family (Crustacea, Arthropoda), all individually selected using a stereomicroscope.

**Table 1:** Amphipods collected by this study for full histopathological screening. Number of individuals belonging to different genera are linked to the location and day of the sampling.

| Sampling location | Location coordinates | Date | Number of amphipods sampled |  |  |  |
| --- | --- | --- | --- | --- | --- | --- |
|  |  |  | *Echinogammarus* sp. | *Gammarus* sp. | *Melita* sp. | *Orchestia* sp. |
| Dart Estuary | 50° 23' 21" N<br>03° 35' 36" W | 19-Sep-16 | 64 | 10 | 10 | 31 |
|  |  | 26-Apr-17 | 40 | 6 | 23 | 21 |
|  |  | 13-Aug-18 | 41 | 0 | 0 | 0 |
| Tamar estuary | 50° 23' 25" N<br>04° 13' 51" W | 20-Sep-16 | 84 | 0 | 4 | 28 |
|  |  | 27-Apr-17 | 41 | 27 | 18 | 37 |
|  |  | 14-Aug-18 | 48 | 0 | 0 | 34 |
|  |  | 08-Nov-18 | 35 | 7 | 0 | 27 |
| Camel Estuary | 50° 32' 17" N<br>04° 56' 05" W | 28-Apr-17 | 30 | 29 | 6 | 37 |
| Newton's Cove<br>(Weymouth) | 50° 36' 17" N<br>02° 27' 03" W | 20-Apr-16 | 50 | 0 | 0 | 0 |
|  |  | 08-Jun-16 | 30 | 0 | 0 | 0 |
|  |  | 13-Sep-16 | 38 | 30 | 13 | 32 |
|  |  | 28-Oct-16 | 30 | 20 | 0 | 0 |
|  |  | 25-Nov-16 | 40 | 0 | 0 | 0 |
|  |  | 14-Dic-16 | 63 | 42 | 9 | 0 |
|  |  | 17-Jan-17 | 40 | 30 | 6 | 0 |
|  |  | 16-Feb-17 | 50 | 23 | 4 | 0 |
|  |  | 16-Mar-17 | 54 | 0 | 0 | 6 |
|  |  | 11-Apr-17 | 38 | 15 | 5 | 0 |
|  |  | 04-May-17 | 51 | 0 | 0 | 0 |
|  |  | 18-May-17 | 31 | 0 | 0 | 0 |
|  |  | 15-Jun-17 | 12 | 10 | 0 | 25 |
|  |  | 21-Jul-17 | 40 | 10 | 0 | 23 |
|  |  | 16-Mar-18 | 55 | 12 | 0 | 10 |
|  |  | 16-Apr-18 | 45 | 8 | 0 | 4 |
|  |  | 11-May-18 | 31 | 0 | 0 | 0 |
|  |  | 13-Jun-18 | 55 | 0 | 0 | 0 |

**Table 2:**
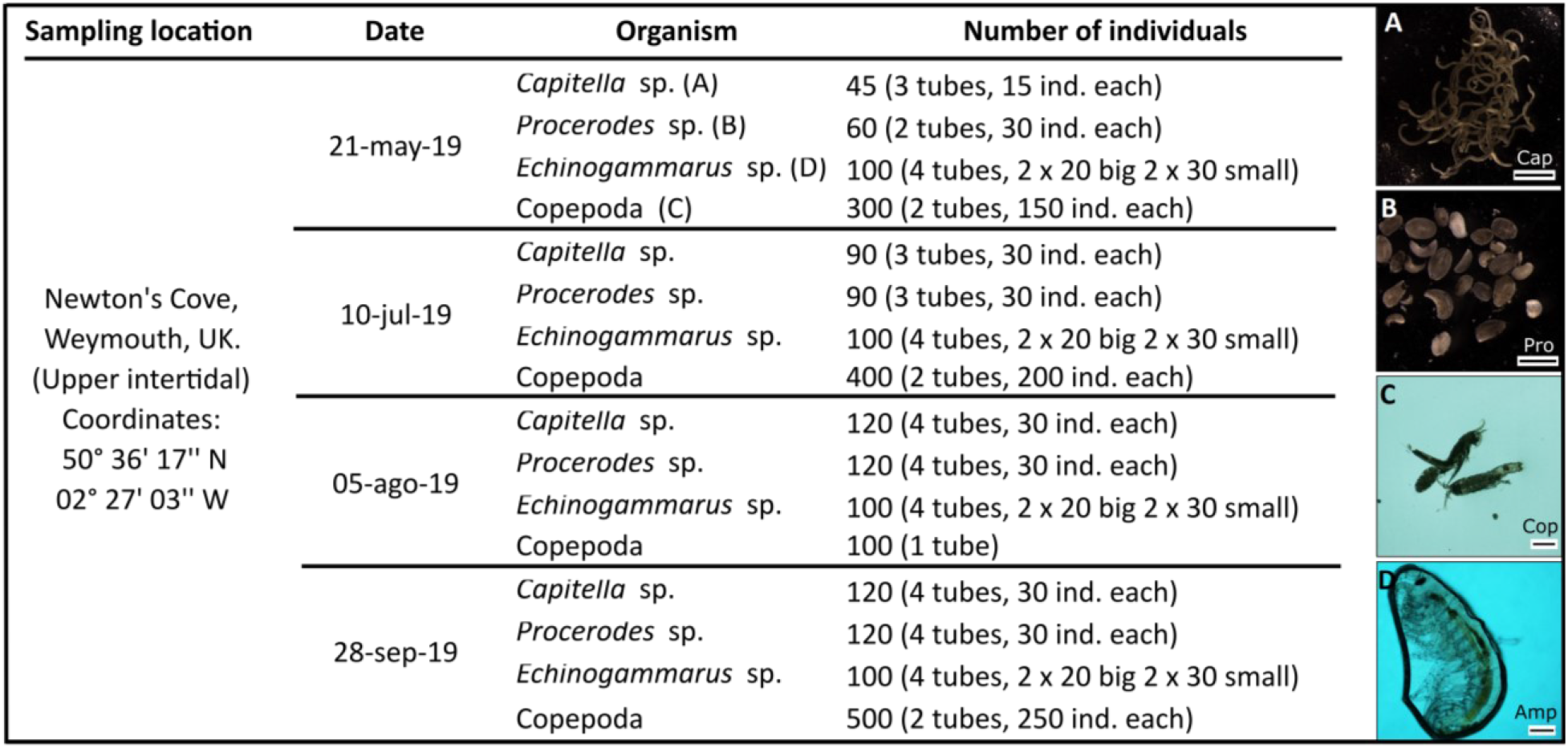
Sampling information for invertebrate species collected for PCR screening of *Txikispora philomaios* in Newton’s Cove. The sampling date, the organism’s genus/clade, and the number of individual organisms included in each batch. Between brackets in “Organism”, stereomicroscopical images of the taxa (A, B, C, D). Between brackets in “No. of Individuals” the total number of individuals per PCR tube.

| Sampling location | Date | Organism | Number of individuals |
| --- | --- | --- | --- |
| Newton's Cove,<br>Weymouth, UK.<br>(Upper intertidal)<br>Coordinates:<br>50° 36' 17" N<br>02° 27' 03" W | 21-may-19 | *Capitella* sp. (A) | 45 (3 tubes, 15 ind. each) |
|  |  | *Procerodes* sp. (B) | 60 (2 tubes, 30 ind. each) |
|  |  | *Echinogammarus* sp. (D) | 100 (4 tubes, 2 x 20 big 2 x 30 small) |
|  |  | Copepoda (C) | 300 (2 tubes, 150 ind. each) |
|  | 10-jul-19 | *Capitella* sp. | 90 (3 tubes, 30 ind. each) |
|  |  | *Procerodes* sp. | 90 (3 tubes, 30 ind. each) |
|  |  | *Echinogammarus* sp. | 100 (4 tubes, 2 x 20 big 2 x 30 small) |
|  |  | Copepoda | 400 (2 tubes, 200 ind. each) |
|  | 05-ago-19 | *Capitella* sp. | 120 (4 tubes, 30 ind. each) |
|  |  | *Procerodes* sp. | 120 (4 tubes, 30 ind. each) |
|  |  | *Echinogammarus* sp. | 100 (4 tubes, 2 x 20 big 2 x 30 small) |
|  |  | Copepoda | 100 (1 tube) |
|  | 28-sep-19 | *Capitella* sp. | 120 (4 tubes, 30 ind. each) |
|  |  | *Procerodes* sp. | 120 (4 tubes, 30 ind. each) |
|  |  | *Echinogammarus* sp. | 100 (4 tubes, 2 x 20 big 2 x 30 small) |
|  |  | Copepoda | 500 (2 tubes, 250 ind. each) |

**Fig 1.**
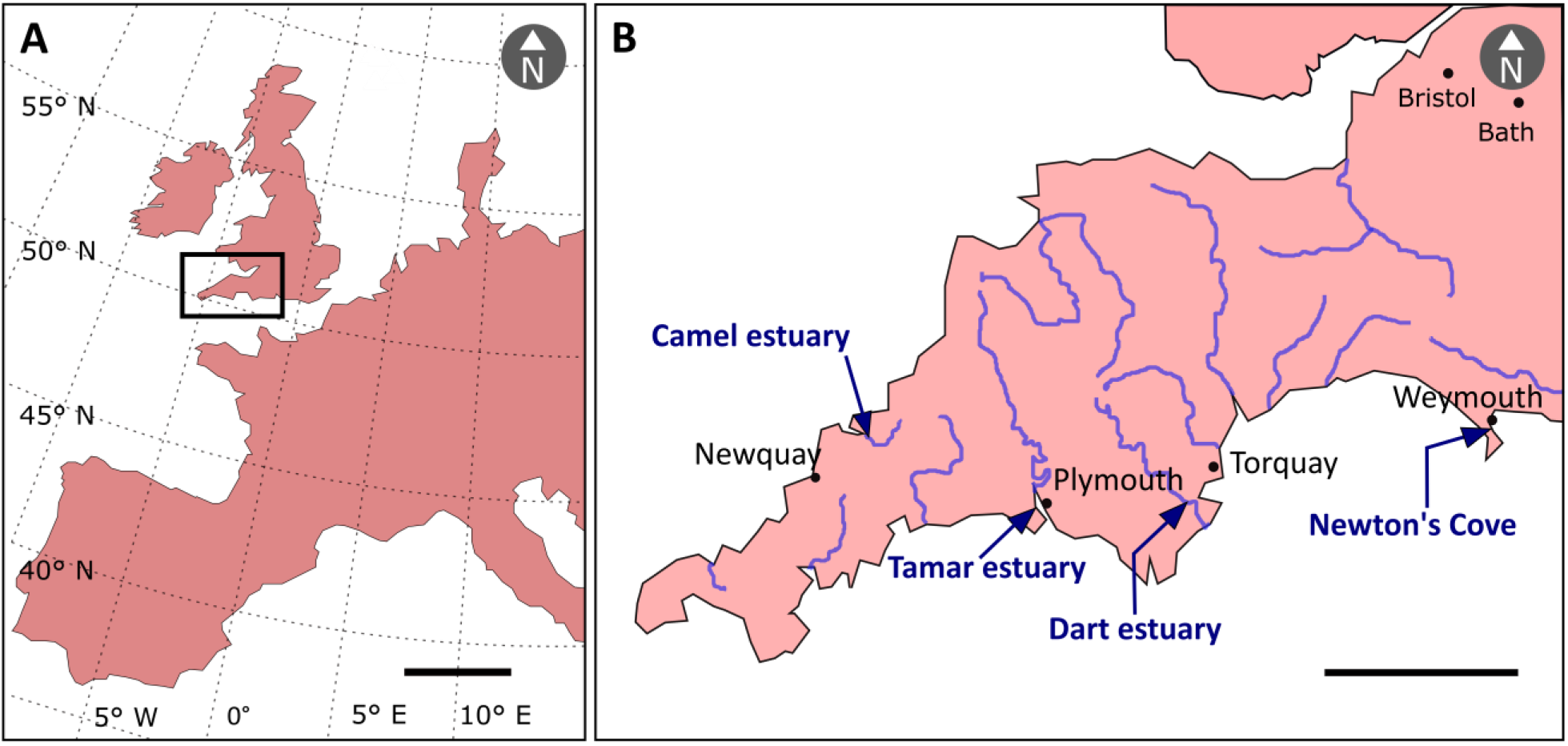
Map showing the coastal locations in which amphipods of the genera *Echinogammarus, Orchestia, Melita* and *Gammarus* were collected. **A**. Western Europe, the black rectangle showing the area of UK sampled. **B**. Area contained within the black rectangle (A). The blue lines show the rivers and estuaries; arrows indicate the sampling locations. Precise coordinates of the locations (Table 1).

### Histology and transmission electron microscopy

Amphipods were kept alive in bottles containing moist algae and dissected within 3-4 hours post collection. The head and two first thoracic segments were fixed in 100% molecular grade ethanol. The following proximate segments of the thorax of about 2 mm in size, were fixed in 2.5% glutaraldehyde in 0.1 M sodium cacodylate buffer (pH 7.4) for TEM. The remainder of the body, which included the last 4-5 segments of the pereon and the pleon, were fixed in Davidson’s seawater fixative (Hopwood 1969) for 24 hours, and then transferred to 70% ethanol. Fresh smears were produced by cutting the distal part of the antennae or uropods before fixation; after a preliminary analysis, slides were left to air-dry. Once dry, slides were stained for 1 minute with Toluidine Blue (1%) and washed with distilled water before being cover-slipped.

For histology, Davidson’s fixed samples were processed from ethanol to wax in a vacuum infiltration processor using established laboratory protocols (Stentiford et al. 2013). Tissue sections (2.5-3 µm) were cut on a Finnese® microtome, left to dry for 24 hours, mounted on VWR^™^ microscope slides, and stained with H&E (Bancroft and Cook 1994). Cover-slipped sections were examined for general histopathology by light microscopy (Nikon Eclipse E800). Digital images and measurements were obtained using the Lucia^™^ Screen Measurement software system (Nikon, UK).

Specimens observed by light microscopy to be infected with *T. philomaios* (one *Echinogammarus* sp. and one *Orchestia* sp.), were selected for TEM analysis. Glutaraldehyde-fixed samples were rinsed in 0.1 M sodium cacodylate buffer (pH 7.4) and post-fixed for 1 hour in 1% osmium tetroxide in 0.1 M sodium cacodylate buffer. Samples were washed in three changes of 0.1 M sodium cacodylate buffer before dehydration through a graded acetone series. Then, they were embedded in epoxy resin 812 (Agar Scientific pre-Mix Kit 812, Agar Scientific, UK) and polymerised overnight at 60 °C. Semi-thin sections (1 µm) were stained with 1% Toluidine Blue and analysed by light microscope, to identify target areas containing sufficient parasites. Ultrathin sections (70-90 nm) were framed on uncoated copper grids and stained with uranyl acetate and Reynold’s lead citrate (Reynolds 1963). Grids were examined using a JEOL JEM 1400 transmission electron microscope and digital images captured using a GATAN Erlangshen ES500W camera and Gatan Digital Micrograph^™^ software.

### DNA extraction, polymerase chain reaction, cloning and sequencing

The head and anterior part of the thorax (preserved in 100% molecular grade ethanol) of 23 amphipods found to be infected via histology (pereon, pleon, and uropods fixed in Davidson’s seawater fixative) were selected for DNA extraction. Infected tissues were disrupted and digested overnight (12 hours) using Fast Prep® Lysing Matrix tubes containing 0.2 mg (6 U) Proteinase K (Sigma-Aldrich®) diluted 1/40 in Lifton’s Buffer (100 mM EDTA, 25 mM Tris-HCl, 1% (v/v) SDS, pH 7.6). Next, a 1/10 (v/v) of 5 M potassium acetate was added to each of the 23 tubes containing digested sample, Proteinase K, and Lifton’s buffer. The solution was mixed and incubated on ice for 1 hour. From here DNA was extracted using the phenol-chloroform method described in (Sambrook et al. 1989). The resulting pellet was diluted in 50 µl of molecular grade water and DNA concentration quantified using NanoDrop^™^ (Thermo Fisher Scientific). *T. philomaios*’ 18S SSU rRNA (hereafter ‘18S’) was amplified by PCR using primers targeting different overlapping regions (Table 3), and the following PCR conditions: A total reaction volume of 20 µl included 10 µl molecular water, 5 µL GoTaq® Flexi Buffer, 2.0 mM MgCl_2_, 0.2 mM of each deoxyribonucleotide, 40 pM of each primer, 0.5 U GoTaq® Polymerase (Promega), and 200 ng of the extracted DNA. The PCR cycling parameters for primer pair (SA1nF + 631R; Bass et al. 2012, and in-house design respectively; Table 3) included denaturation for 5 minutes at 95 °C, followed by 35 cycles alternating: 95 °C (30 s), 57 °C (30 s), and 72 °C (90 s); before a final extension and incubation of the amplicons at 72 °C for 10 minutes. Same conditions were used for primer combinations (S47-152F + S47-617R and S47-472F + S47-1027R; Table 3) except for the annealing temperature which was 67 °C (30 s). Amplicons were cleaned using 20% polyethylene glycol 8000 (Sigma-Aldrich®) followed by ethanol precipitation, and a-tailed to improve cloning efficiency before another PEG 8000 clean. Clone libraries were created using Strategene’s cloning kit (Agilent Technologies, Santa Clara, CA, USA) as per manufacturer’s protocol. Bacterial colonies were picked from LB/ampicillin plates and suspended in 20 µl PCR water and lysed at 95 °C for 5 minutes. Eight clones from each library were amplified with 1 µl lysed culture DNA and M13F/M13R primers (Invitrogen^™^– Thermo Fisher Scientific) using the mastermix concentrations described previously, and the manufacturers program. PCR products were bead-cleaned and a total volume of 15 µl was mixed with 2 µl of the M13F forward primer, before being single-read Sanger sequenced (Eurofins^®^Genomics).

**Table 3:**
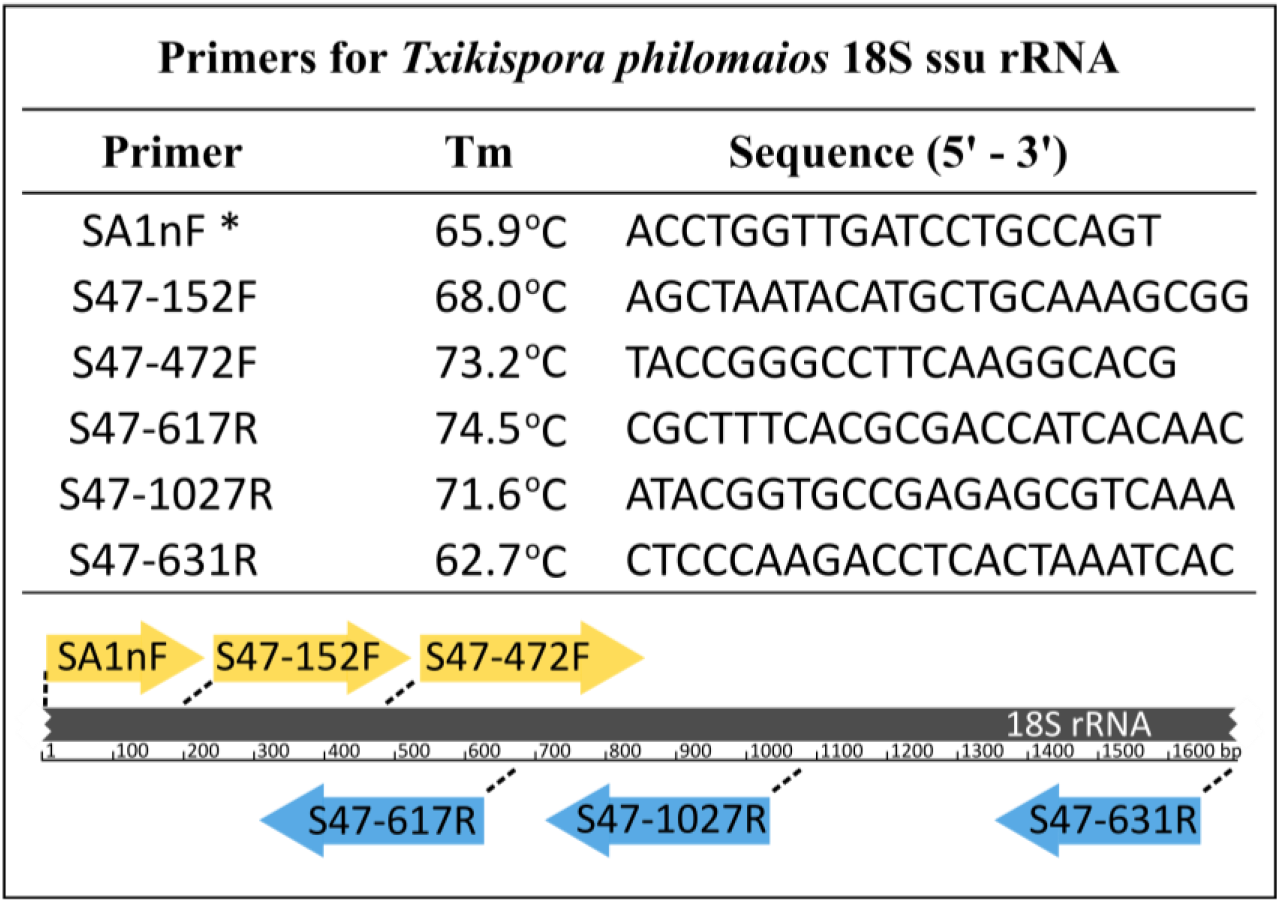
List of primers designed for *Txikispora philomaios* amplification and universal primer SA1nF (*) from Bass et al. 2012. The melting temperature (Tm) and the sequence for each primer is specified. In the bottom, a diagram indicating position of attachment for each primer in the 18S ssu rRNA and direction of amplification.

### *In-situ* hybridization

Tissue sections (4 µm) from the individuals of interest were recovered from the 42 °C water bath (without Sta-On tissue-adhesive) using Polysine® Slides (Thermo Fisher Scientific) and left to dry for 24 hours. The forward S47-152F and reverse S47-617R primers were used to amplify part of the 18S extracted from an infected individual of *Orchestia* sp. DNA amplification and purification were carried out using the same concentrations and conditions explained in previous section. Purified DNA was digoxigenin (DIG)-labelled using same primers and PCR conditions above, but changing the concentration of reagents, say: 10 μl 5X Colorless GoTaq® Reaction Buffer, 5 μl MgCl_2_ solution (Promega), 5 μl of PCR DIG labelling mix (Roche), 3 μl template DNA, 1 μl of forward and reverse primers, 0.5 μl of GoTaq Polymerase, and 24.5 μl molecular grade water. The control slide was produced amplifying the same 18S region using non-labelled standard DNTPs. Products generated via PCR were purified as described in previous section, total DNA quantified (NanoDrop 1000 Spectrophotometer® Thermo Scientific) and diluted to 1 ng/μl for a total volume of 50 μl.

Dry tissue sections were dewaxed and rehydrated: Clearene for 5 minutes (2 times), followed by 100% IDA (industrial denatured alcohol) for 5 minutes and 70% IDA another 5 minutes. Slides were rinsed in 0.1M TRIS buffer (0.1 M TRIS base, 0.15 M NaCl, adjust the pH to 7.5 adding HCl) and placed in a humid chamber. Each slide was covered with 300 μl of 0.3% Triton-X diluted in 0.1M TRIS buffer (pH 7.5) for 20 minutes and rinsed with 0.1M TRIS buffer (pH 7.5). Tissue was covered with Proteinase K diluted to 25 μg/ml in prewarmed (37 °C) 0.1M TRIS buffer (pH 7.5) and kept for 20 minutes at 37 °C within the humid chamber to prevent evaporation. Slides were washed in 70% IDA for 3 minutes and 100% IDA for another 3 minutes before rinsing them in SSC 2X for 1 minute while gently agitating (SSC 1X is 0.15 M sodium chloride and 0.015 M sodium citrate). Slides were kept in 0.1 M TRIS buffer (pH 7.5) until the *in-situ* hybridization frame seals (BIO-RAD) were glued to the slide around the sample. Then, the DIG-labelled probe and the non-labelled probe (control), both 50 µl in volume, were diluted by adding 50 µl of hybridization buffer and added to the cavity created by the gel frames in the slide, with the sample in the middle. After DNA denaturation at 94 °C for 6 minutes, slides were hybridized overnight (16 h) at 44 °C.

Samples were washed for 10 minutes with room temperature washing buffer (25 ml of SSC 20X, 6M Urea, 2 mg/l BSA), before being washed twice with preheated (38 °C) washing buffer for 10 minutes each. Slides were rinsed with preheated (38 °C) SSC 1X for 5 minutes (2 times) and with 0.1M TRIS buffer (pH 7.5) another 2 times. The blocking step was carried out with a solution of 6% dried skimmed milk diluted in 0.1M TRIS buffer (pH 7.5) for 1 hour at room temperature and washed with 0.1M TRIS buffer (pH 7.5) for 5 minutes twice.

Slides were incubated with 1.5 U/ml of anti-DIG-AP Fab fragments (Roche) diluted in 0.1M TRIS buffer (pH 7.5) for 1 hour at room temperature in darkness. The excess of Anti-DIG-AP was removed by 4 successive washes in 0.1M TRIS buffer (pH 7.5) for 10 minutes each. Slides were transferred to 0.1M TRIS buffer (pH 9.5) which is (0.1M TRIS base, 0.1M NaCl, adjust pH to 9.5 adding HCl) for 2 minutes and then tissue was covered with NBT/BCIP stock solution (Roche) diluted in 0.1M TRIS buffer (pH 9.5) at 20 μl/ml, and incubated in darkness and room temperature until the first clear signs of blue staining appeared (about 30 minutes). Slides were washed in 0.1M TRIS buffer (pH 9.5) for 1 minute twice and stained with 1% Bismark Brown for 6 minutes. Finally, slides were dehydrated by immersing them for 30 seconds in 70% IDA, 45 seconds in 100% IDA and 2 washes in clearene for 1 minute each. Slides were air dried for 30 minutes and permanently cover-slipped with DPX mounting medium (Sigma-Aldrich).

### Sequence alignment and phylogenetic analysis

The PCR-amplified 18S rRNA was BlastN-searched (Zhang et al. 2000) against the GenBank nucleotide (nt) database. Holozoan 18S rDNA gene sequences, as well as sequences from those uncultured organisms showing highest similarity, were downloaded and aligned with the consensus 18S rDNA gene sequence from *T. philomaios* in MAFFT v.7 (Katoh et al. 2017) using the accurate option L-INS-i. The alignment was trimmed by trimAl v.1.4.rev22 (Capella-Gutiérrez et al. 2009) using the (-gt 0.1) option, and manually curated in SeaView v.4 (Gouy et al. 2010). In turn, the best-fitting model (GTR + F + G) for the alignment was selected using ModelFinder (Kalyaanamoorthy et al. 2017) as implemented in IQ-TREE v.1.6.10 (Nguyen et al. 2015) and used to generate a ML tree in IQ-TREE. Branch support was obtained from 1000 ultrafast bootstrap values (Minh et al 2013). A second maximum likelihood phylogenetic tree was constructed using RAxML v8.2.12 (Stamatakis 2014); support values calculated using 1,000 bootstrap replicates were mapped onto the tree with the highest likelihood value (evaluated under GTRGAMMA model). A Bayesian inference consensus tree was built using MrBayes v.3.2 (Ronquist et al. 2012) under default parameters except for the following: the number of substitution types was mixed; the model for among-site rate variation, Invgamma; the use of covarion like model, activated. The MCMC parameters changed were: 5 million generations; sampling frequency set to every 1,000 generations; burnin fraction value = 0.25; starting tree set to random, and all compatible groups consensus tree. A final consensus tree figure was created using FigTree v1.4.3 (Rambaut 2017) and based on the Bayesian topology.

A second 18S phylogenetic tree was constructed including environmental and unclassified sequences branching with or within Filasterea, by mining different databases. The 18S rDNA gene of *T. philomaios* was used as a bait to fish highly-similar sequences, by blastn searching against the following GenBank archives: nt, whole genome shotgun contigs (WGS), sequence read archive (SRA), and high throughput genomic sequence archive (HTGS). The same approach was followed for SILVA (www.arb-silva.de), ENA (www.ebi.ac.uk) and DDBJ (www.ddbj.nig.ac.jp) databases. All environmental sequences branching within Filasterea or sister to it in a preliminary tree were retained for subsequent analyses (Table 4), as were as a selection of highly divergent uncultured mesomycetozoean and choanoflagellate sequences. Sequences belonging to uncultured organisms that branched robustly to existing species in Ichthyosporea, Choanoflagellatea or Metazoa were excluded from the final alignment (the selected sequences were realigned). The alignment and subsequent phylogenetic analysis were constructed as described above.

**Table 4:**
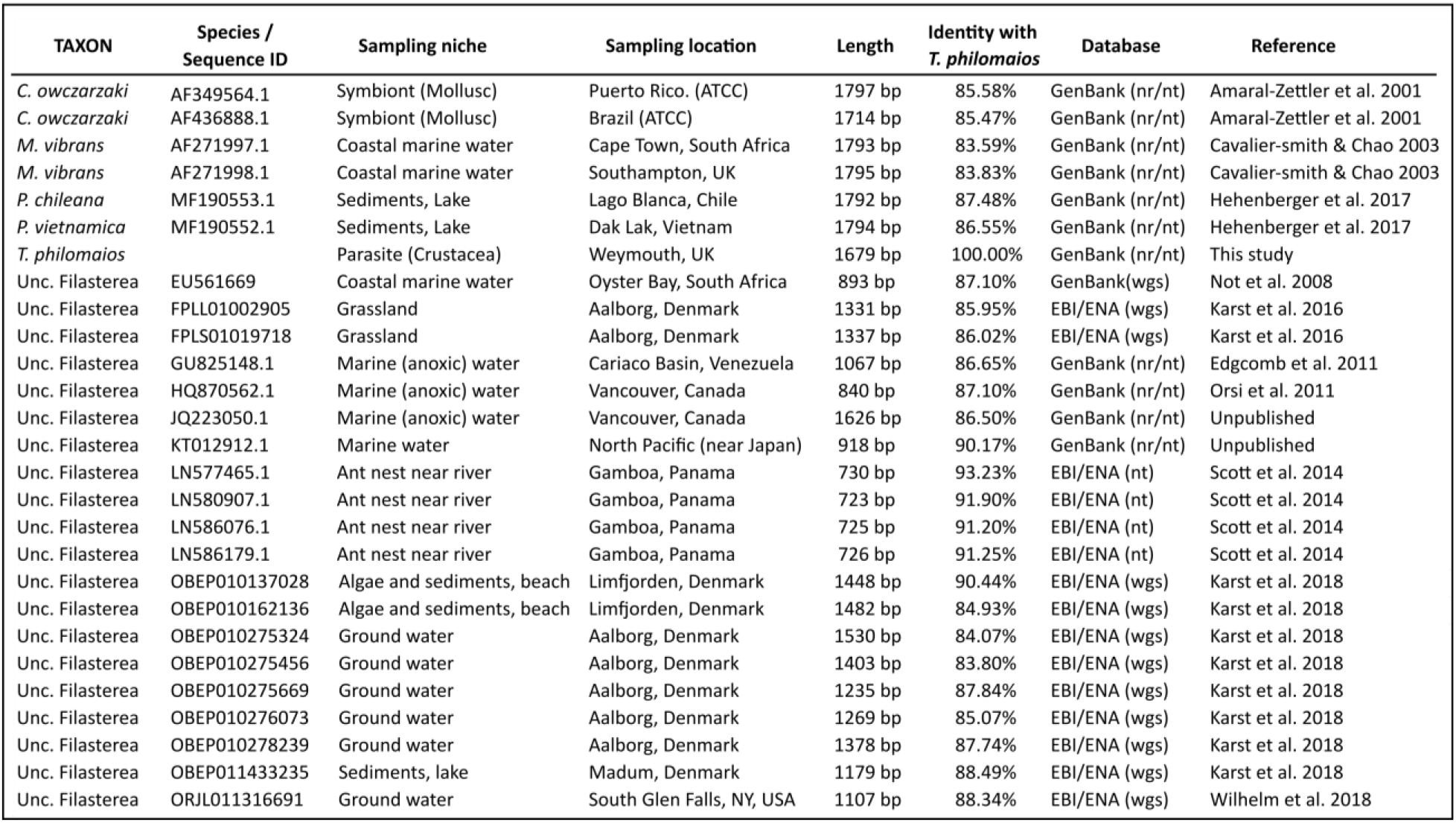
List of existing filastereans and uncultured organisms associated to this lineage according to our phylogenetic analysis (Figs. 10, 11). The sequence ID corresponds the name used in the phylogenetic trees, and it is linked to the ecosystem (sampling niche) and the geographic site (sampling location) from which the 18S ssu rRNA was collected. In the case of parasites and symbionts, susceptible hosts have been specified as the sampling niche. The list also includes the sequences’ length, its identity (percentage) to *T. philomaios’* 18s, the database from which it was mined, and the reference to the authors who uploaded/published it.

## RESULTS

### Clinical signs and prevalence

Two amphipod genera, *Echinogammarus* and *Orchestia*, were found infected by *T. philomaios*. The genera *Gammarus* (n = 279) and *Melita* (n = 101) were also investigated, but no signs of infection were observed histologically. However, the number of individuals examined was considerably lower (Table 1). Infection by *T. philomaios* was suggested macroscopically in heavily infected individuals by a yellowish and opaque tegument (Fig. 2). The carapace thickened and lost rigidity (Fig. 2B), impeding to discern internal organs, especially the intestine, which is evident in young healthy individuals. Besides, gross examination of the most translucent appendages (antennae, uropods, and gills) using a stereomicroscope permitted detection of the parasite in haemolymph (Fig. 3). Infected individuals displayed lethargy, unresponsiveness to stimuli, and very reduced jumping ability in the case of the sandhopper (*Orchestia* sp.).

**Fig 2.**
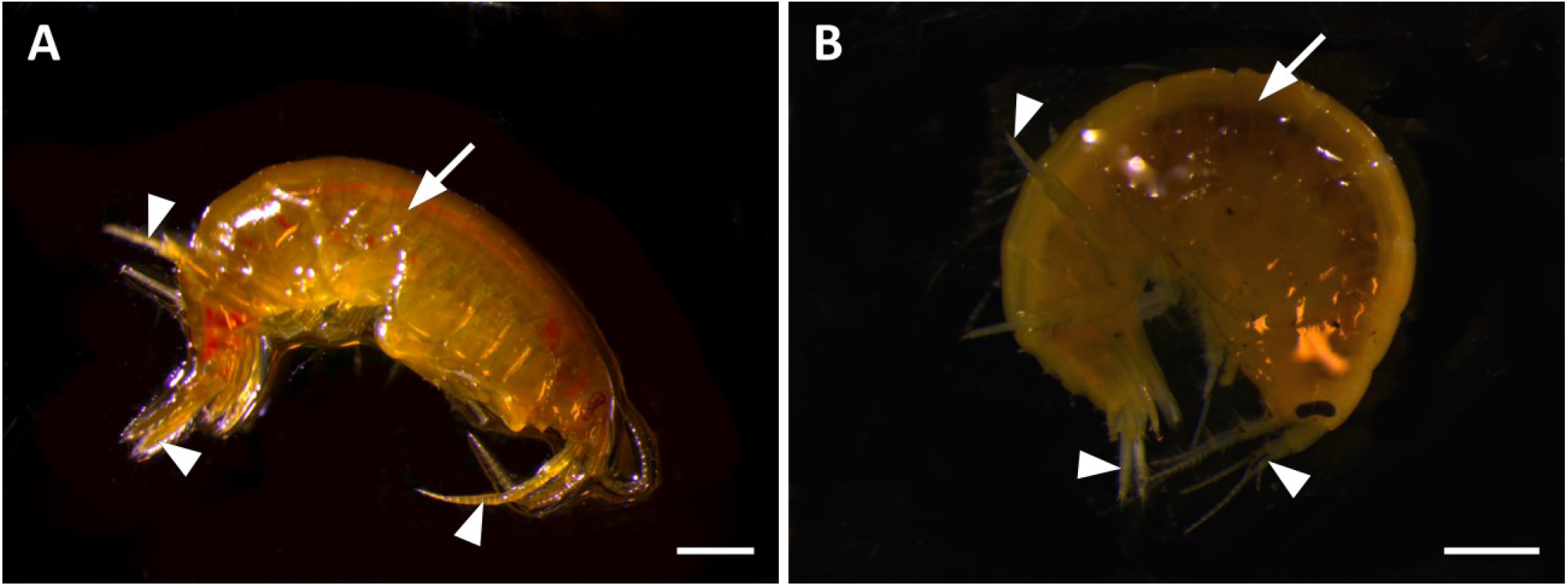
Stereo-microscopical images of live *Echinogammarus* sp. amphipods collected in Newton’s Cove. **A**. Uninfected individual. Antennae, pereopods and uropods (arrowheads), internal organs (arrow) **B**. Individual heavily infected by *Txikispora philomaios*. The tegument of the infected individual appears more opaque, the gut (arrow) is not evident, especially in the posterior fraction of the body (pleon). Scale bars = 100 µm for (A & B).

**Fig 3.**
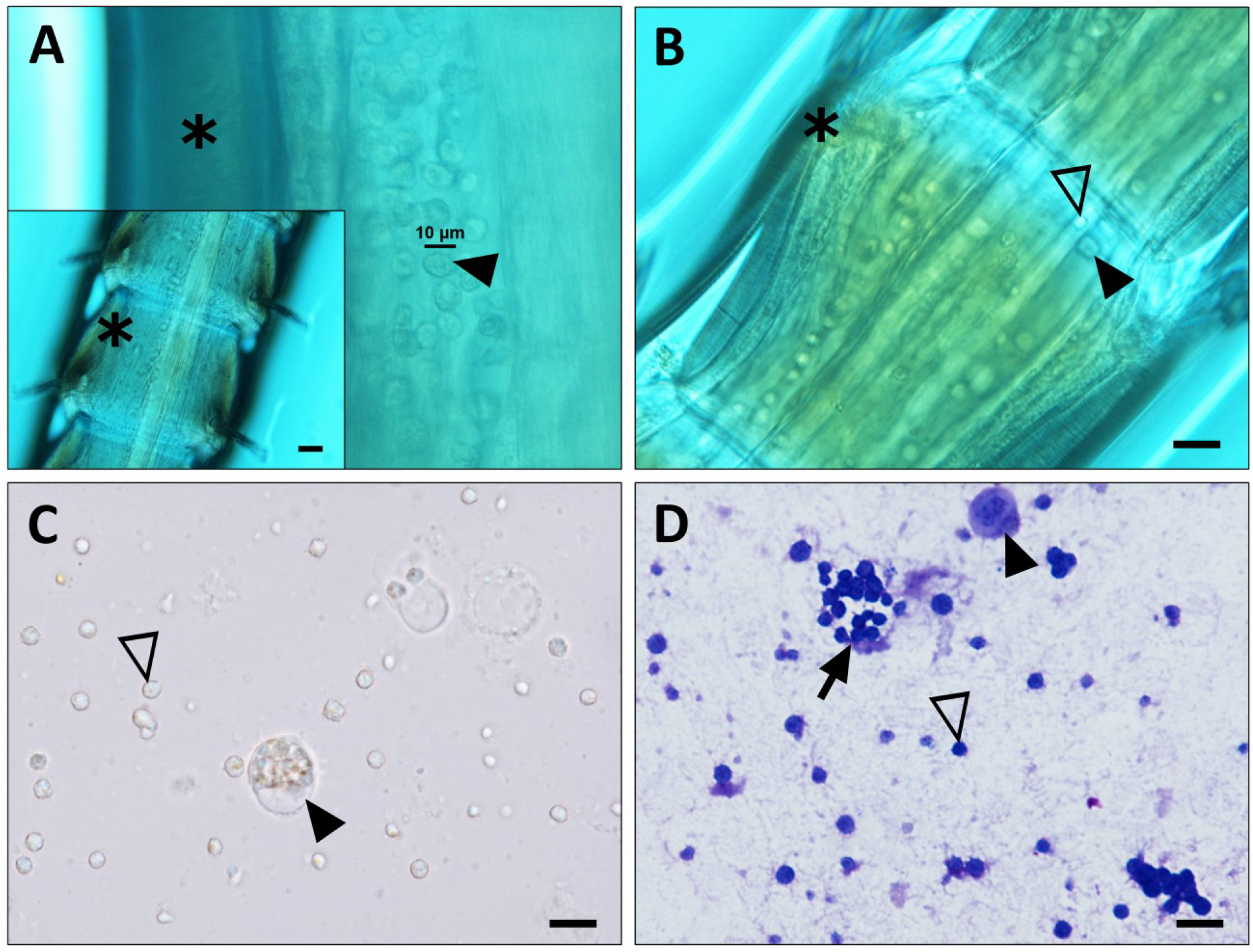
Light microscopic images of antennae (A, B), and haemolymph (C, D) from healthy (A) and infected (B, C, D) amphipods of genus *Echinogammarus*. **A**. Stereo microscope image of the antennae (inset) of a healthy amphipod individual, showing (≈ 10 µm) haemocytes (arrowhead) flowing in the open circulatory system between the antennal gland and the tegument (asterisk). **B**. Cells of *Txikispora philomaios* (empty arrow) can be differentiated from haemocytes (filled arrowhead) by their smaller size and small nucleus. **C**. Composed microscope image of an unstained fresh preparation of the haemolymph showing *T. philomaios* cells free in the haemolymph (empty arrowhead) and within haemocytes (filled arrowhead). **D**. Toluidine blue-stained preparation of haemolymph from an infected amphipod showing *T. philomaios* single cells (empty arrowhead), parasitic cells inside haemocytes (filled arrowhead), and parasite cells forming multicellular groups (arrow). Scale bars = 10 µm for (A, B, C, D), and 20 µm for inset in (A).

Discrimination between haemolymph cells (8-10 µm) and *T. philomaios* cells (2-4 µm) was possible on the basis of the cell diameter and nuclear size (Fig. 3A, 3B). Haemolymph smears (Fig. 3C) evidenced the difference between the spherical and peripheral nucleus of *T. philomaios* (∼1 µm) and the central and irregular one in haemocytes (6-8 µm) (Fig. 3C). Additionally, fresh preparations allowed to notice the occurrence of up to 10 parasite cells inside hosts haemocytes. Toluidine staining of the dry smears emphasised the structures, allowing the observation of cell aggregates (Fig. 3D). The occurrence of *T. philomaios* infection was consistent throughout the years of study (2016-2018) showing a distinct prevalence peak between late April and early June; at least for the regularly sampled *Echinogammarus* sp. population present in Newton’s Cove. These outbreaks of *T. philomaios* infection were usually short-lived, usually lasting no more than three weeks. However, the prevalence of infection was high, varying between 24% (2018) and 64% (2016) in the coastal location of Weymouth. Although the limited data from the other sampling sites precluded direct comparison, the parasite was present in the Dart, Tamar and Camel estuaries at low levels in both spring and autumn (Fig. 4A). *Orchestia* sp. was less frequently and abundantly sampled, but in Newton’s Cove, infection also seemed to peak during May and early June (Fig. 4B). While in *Orchestia* sp. sampled in Newton’s Cove the prevalence was lower (10%), the parasite was more frequently detected in the Dart and Tamar estuaries. The prevalence of infections in *Echinogammarus* sp. during the rest of the year (from June to early April) was low (1.9%, n = 1136), and infection was never systemic. The few parasitic cells observable during these months were almost exclusively associated with the testis.

**Fig 4.**
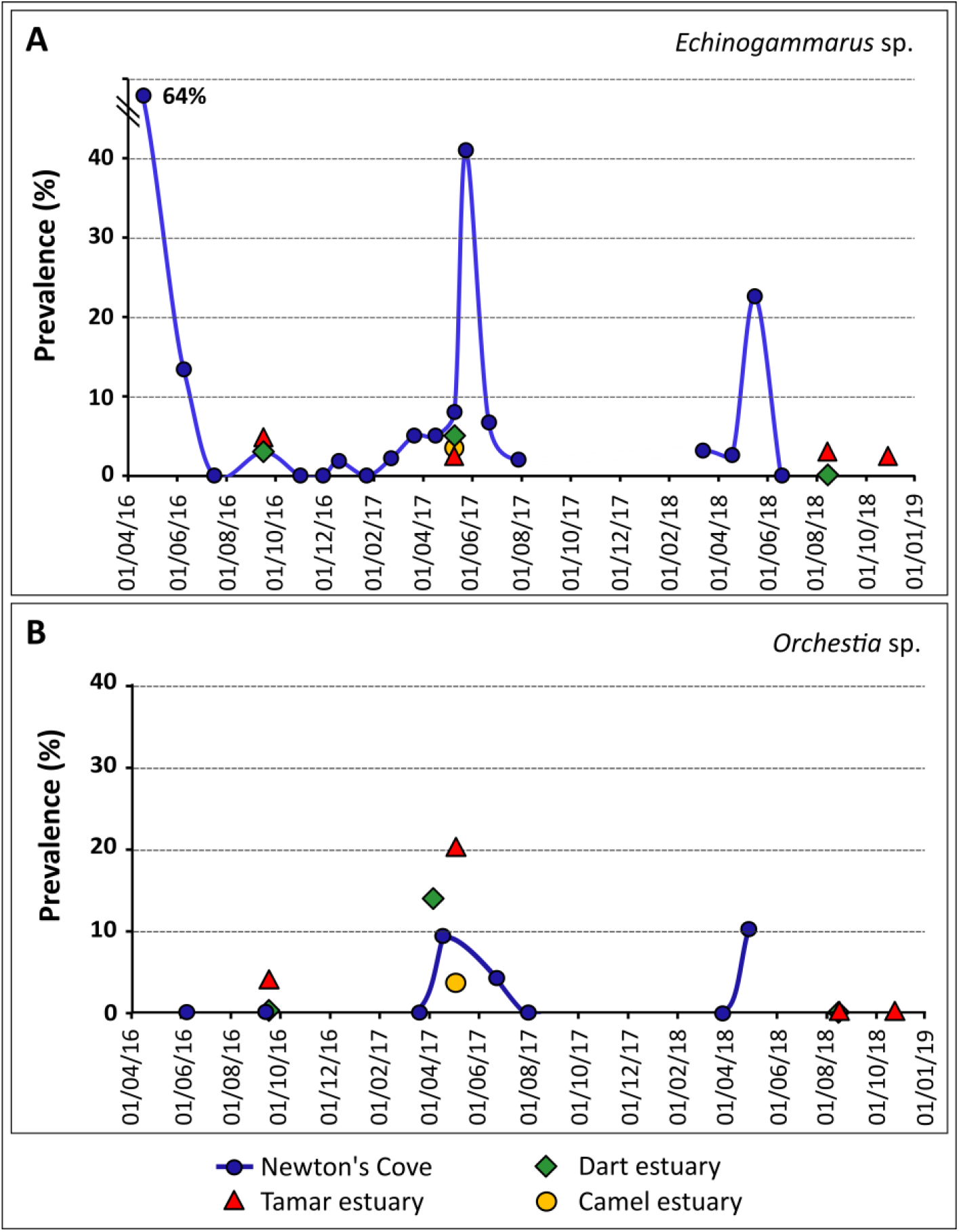
Prevalence of *Txikispora philomaios* infection in *Echinogammarus* sp. (**A**.), and *Orchestia* sp. (**B**.) from April 2016 to August 2018. Dates on the x-axis correspond to sampling information in (Table 1, 2). Y-axis: *T. philomaios* infection prevalence (%). Blue spheres refer to amphipods collected in Newton’s Cove; red triangles = Tamar estuary; green diamonds = Dart estuary; yellow spheres = Camel estuary.

### Histopathology and ultrastructure

Cells of *T. philomaios* were virtually spherical (width = 1.94 ± 0.21 µm; length = 2.36 ± 0.23 µm; n = 50) when fixed in Davidson’s seawater fixative, and 2.26 ± 0.34 µm by 2.60 ± 0.41 µm; n = 50) when preserved in glutaraldehyde. By light microscopy, a nucleus in the periphery of the cell was distinguished in a very translucent cytoplasm. Parasites were present in the haemolymph and frequently intracellularly within haemocytes (Fig. 5A). Infected haemocytes (containing up to 10 *T. philomaios* cells) were often necrotic, with a clear loss of cellular integrity. In contrast, the parasites inside them appeared to be intact. Aggregates of *T. philomaios* cells occurred free or within haemocytes, where similar sized stages were contained within a membrane. However, it was not possible to discern by light microscopy if aggregation was the result of single cells actively joining, or clusters of cells remaining together after the rupture of the haemocyte containing them. Proliferation of *T. philomaios* cells was associated with congestion of haemal sinuses of the tegumental gland and the connective tissue associated with the cuticular epithelium (Fig. 5B). In such systemic infections (with haemolymph, connective and tegument affected), *T. philomaios* was frequently observed infecting the hepatopancreas (Fig. 5C), and seldom in nervous tissue. In the hepatopancreas, the parasite was associated with structural damage with significant inflammation and granuloma formation, often encapsulating *T. philomaios* cells (Fig. 5C). The testis and ovary (Fig. 5D, 5E) also became infected, notably in early-stage infections that did not show evidence of the parasite in other organs and tissues. However, intracellular infections in oocytes and spermatozoids were not observed.

**Fig 5.**
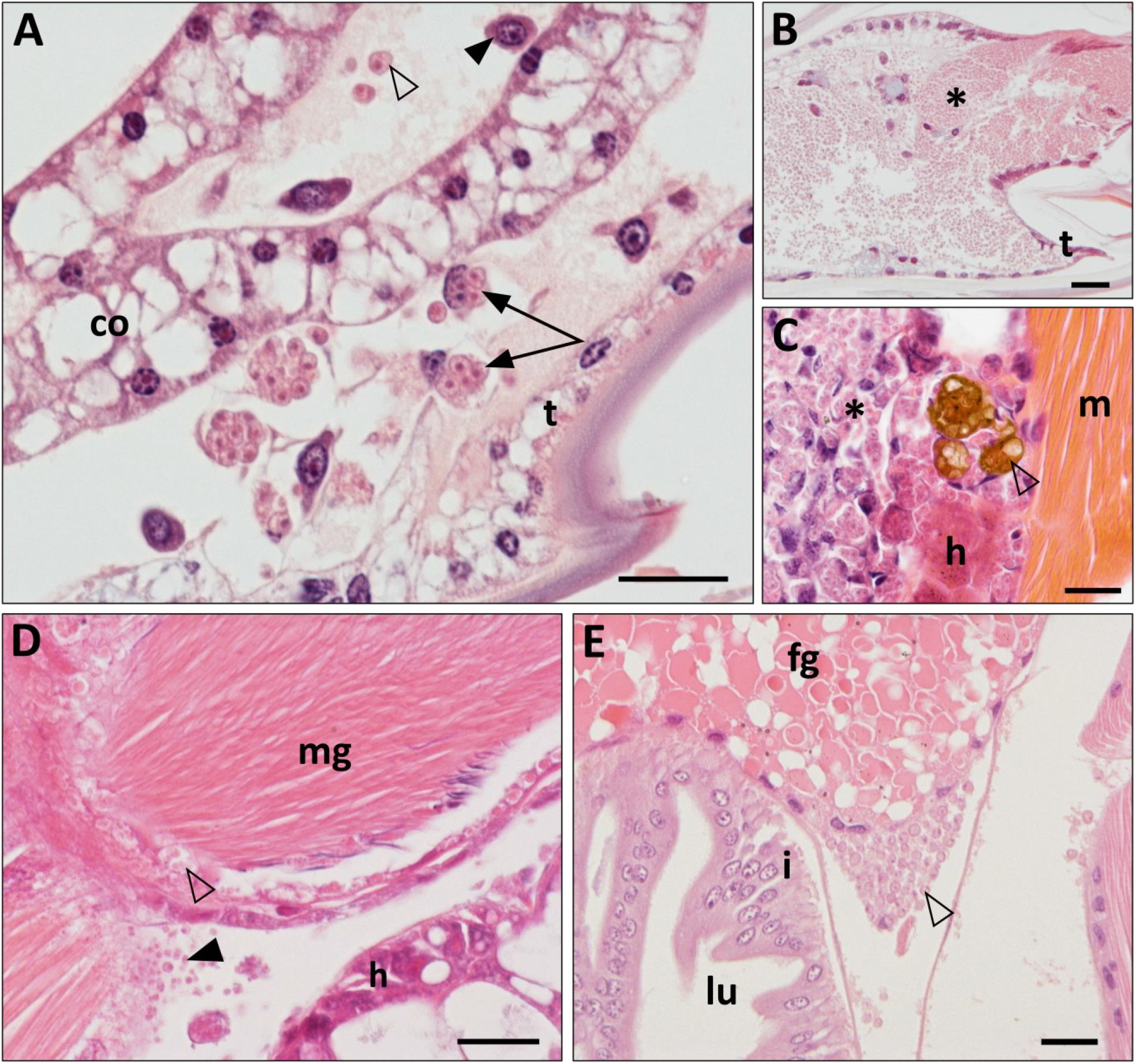
Histological appearance of *Txikispora philomaios* infecting different tissues in *Orchestia* sp. **A**. Parasite cells were observed free in the haemolymph (empty arrowhead) and inside haemocytes (arrows). Non-infected haemocytes (filled arrowhead), tegument (t) and connective tissue (co), in the pereopods of the amphipod. **B**. Masses of parasitic cells (*) in the haemolymph and tegumental gland (t) associated to the cuticle of the carapace. **C**. Parasite cells (*) infiltrating the hepatopancreas (h). Granulomas and melanization (empty arrowhead) and muscle fibres (m). **D**. *T. philomaios* cells infiltrated between muscle fibres (filled arrowhead) and inner connective layers (empty arrowhead) of male gonads (mg). **E**. Disrupted female gonadal tissue (fg) associated to parasitic cells (empty arrowhead). Unaffected intestine (i) and its lumen (lu). Scale bars = 20 µm for (A, B, C, D, E).

At the TEM, *T. philomaios* was found more often as single cells, but also forming clusters containing 3-4 cells (Fig. 6A, 6B). Single cells, often coated by a cell wall, contained a pale staining nucleus with a peripheral compact nucleolus, small mitochondria with lamellar cristae and lipoid structures of varying electron-density (Fig. 6A). These lipoid inclusions displayed morphologic plasticity and variable staining characteristics between *T. philomaios* cells of different size. (Fig. 6C, 6D). Electron-lucent granules appeared integrated within the cytoplasm, while darker granules were often membrane bound and associated to evaginations of the cell wall (Fig. 6G, 6H). The multi-layered cell wall varied in thickness (Fig. 6G, 6H, 6I) and in approximately 30% of the cells examined, appeared detached from the plasma membrane (Fig. 6A, 6G). In few cases, a matrix was observable between cell wall and the detached plasma membrane (Fig. 6C).

**Fig 6.**
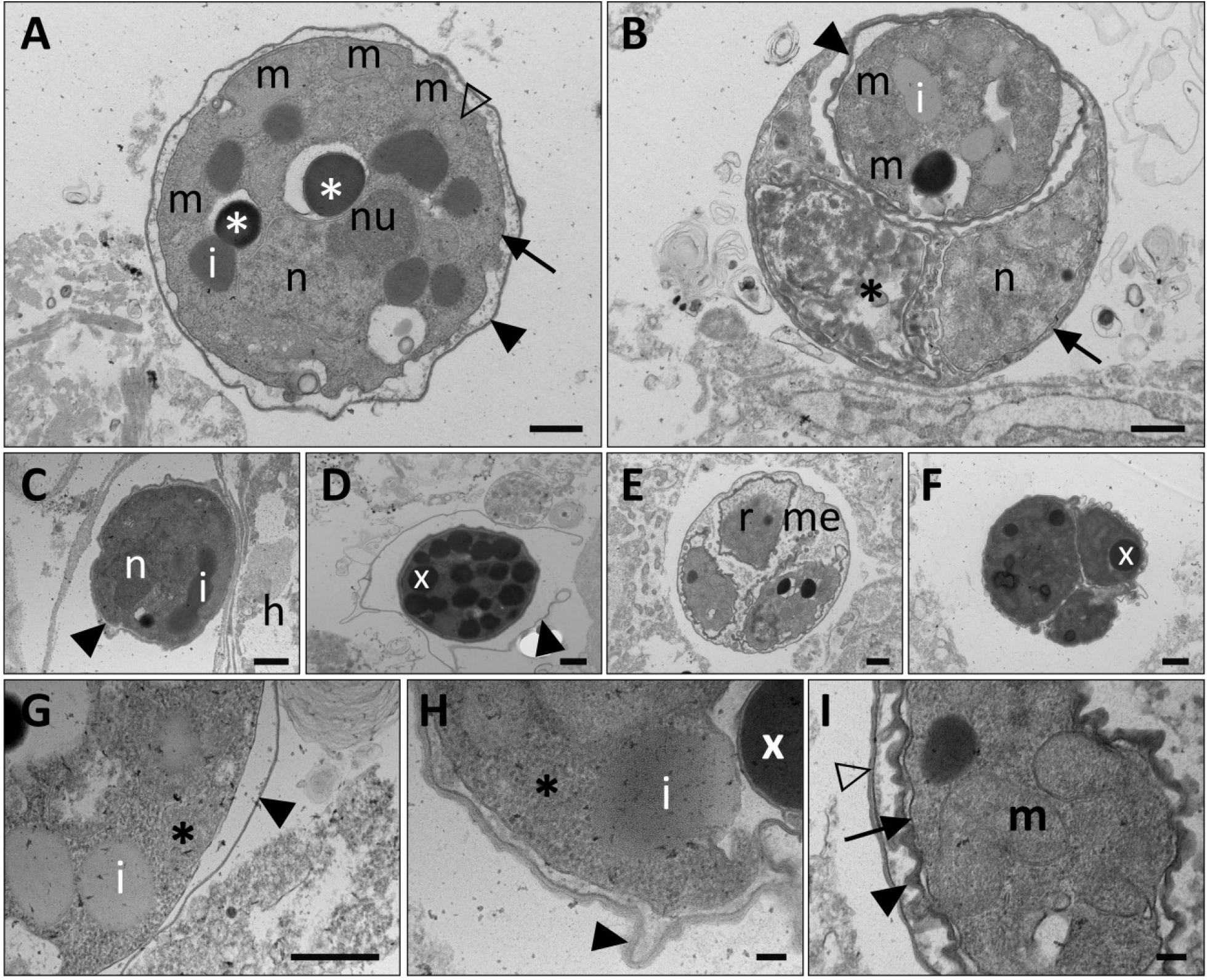
Transmission Electron Microscope (TEM) micrographs of *Txikispora philomaios* cells infecting *Orchestia* sp. **A**. Unicellular stages of the parasite show a single amorphous nucleus (n) with a peripheral nucleolus (nu), peripheric mitochondria (m) electron-dense lipidic vesicles (*), and electron-lucent vesicles (i). The cell wall (filled arrowhead) appears detached from the plasma membrane (arrow). **B**. Dividing form of the parasite, with outer cell wall (arrow) and walled inner cells (filled arrowhead). One of the inner cells appears necrotic (*). **C**. Unicellular stage attached to host cell (h); amorphous material between wall and plasma membrane (filled arrowhead); (i) electron-lucent vesicles (reserve material). **D**. Unicellular stage full of electron-dense vesicles (x) with disrupted cell wall around (filled arrowhead). **E**. Dividing form, with inner cells (r) partially sharing the same matrixial material (me) with the outer walled cell. **F**. Electron-dense tricellular stage still within an indistinct walled outer cell. **G**. Detail of the thin wall (filled arrowhead) of a unicellular parasite cell inside a host haemocyte. Electron-lucent vesicles (i) and granular cytoplasm (*). **H**. Detail of a unicellular parasite cell with a thickening and evaginating cell wall (arrowhead). **I**. Detail of outer (empty arrowhead) and inner (filled arrowhead) cell walls, plasma membrane (arrow), and mitochondria (m). Scale bars = 500 nm for (A, B, C, D, E, F, G) and 100 nm for (H, I).

A multicellular stage of *T. philomaios* was also frequently prominent (Fig. 6B); tricellular in appearance a hidden fourth cell was occasionally observed (Fig. 7D). In several multicellular clusters (Fig. 6B, 6E, 6F) the cells were indistinguishable from the unicellular stages present in the haemolymph (Fig. 6A, 6C, 6D). Occasionally, one or more individual cells contained within the walled parent cell were necrotic (Fig. 6B). Numerous peripheral mitochondria were observed in cells with a thickened wall (Fig. 6A, 6I). The thickening of the electron-dense wall of the inner cells was concurrent with a diminishing wall of the receptacle (Fig. 6D, 6F). The presence of unicellular and divisional forms of *T. philomaios* inside host haemocytes and tegumental gland hinted by light microscopy was corroborated by TEM analysis (Fig. 7A, 7B, 7C). Parasite cells appeared healthy in contrast to the compromised integrity of the infected host cell (Fig. 7C). The multicellular form appeared more often within haemocytes (Fig. 7A, 7B), while unicellular stages were more commonly observed free in the haemolymph or inside cells of the host tegument (Fig. 7C)

**Fig 7.**
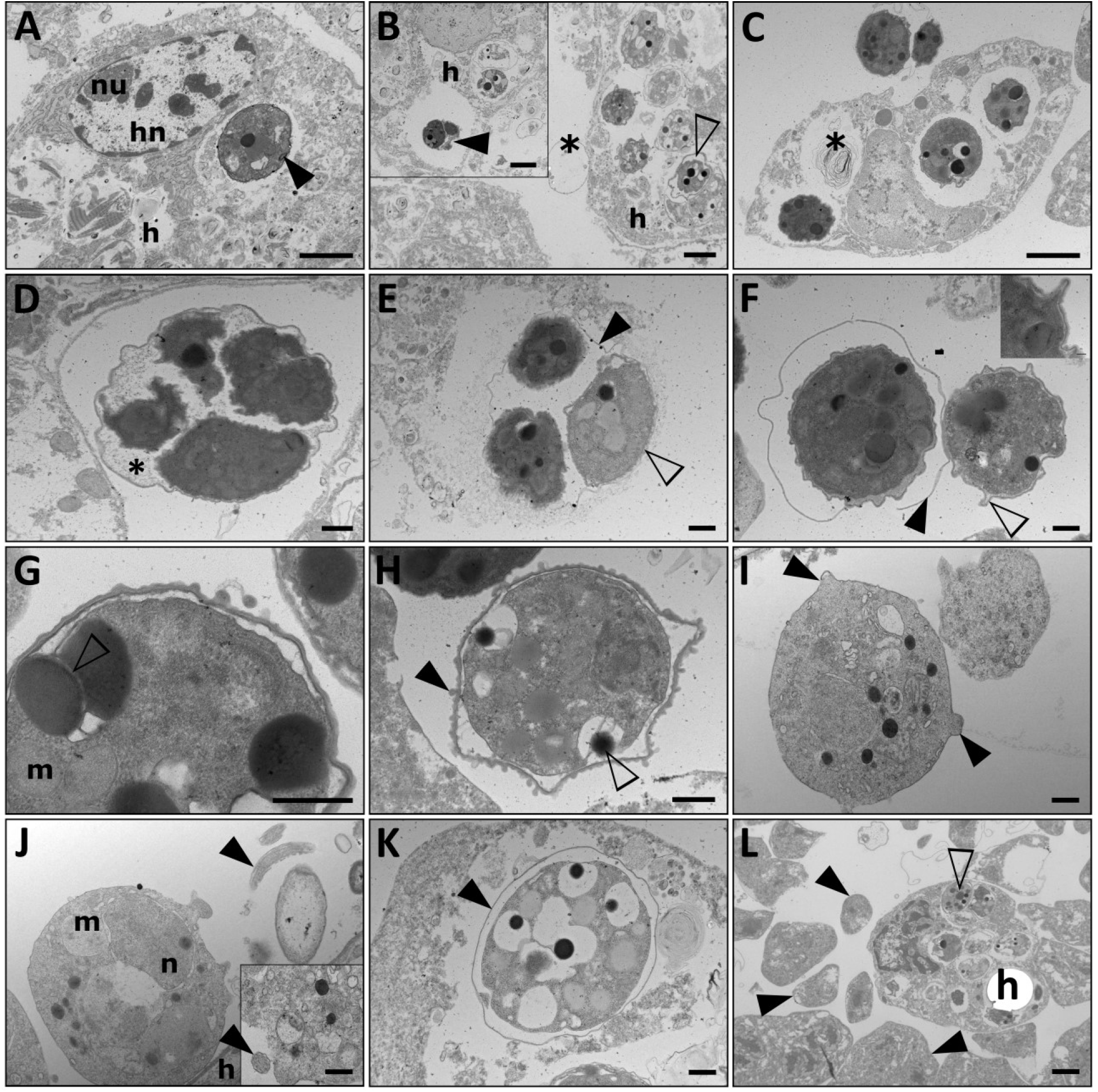
Transmission Electron Microscope (TEM) micrographs of *Txikispora philomaios* cells infecting *Orchestia* sp. **A**. Intracellular stage of *T. philomaios* with a fine and closely attached cell wall (filled arrowhead). The host (h), its nucleus (hn), and nucleolus (nu) are shown. **B**. Five unicellular and a single multicellular stage (empty arrowhead) inside a host haemocyte (h), with a presumed parasite cell wall (*) attached to it. The inset shows the presence of a more electron-dense dividing form of *T. philomaios* (filled arrowhead) inside a host cell (h). **C**. Necrotic haemocyte containing three intact *T. philomaios* cells, with one vacuole containing a necrotic *T. philomaios* cell (*). The infected host cell is unable to maintain its normal structure, also true for its nucleus (hn). **D**. Divisional stage of *T. philomaios*. Four electron-dense daughter cells increase in size inside the wall of the parent cell, which still contains an evident cytoplasmatic matrix (*). **E**. Three daughter cells inside a parent cell without matrix and a very reduced cell wall (arrowhead). One of the daughter cells is more translucent (empty arrowhead) than its sister cyst-like cells. **F**. Two unicellular stages, one of them with an open thin wall (filled arrowhead) similar to the one marked with an asterisk in figure 7B. The other with short projections of the outer cell wall (empty arrowhead). Detail of the inner structure of the projection in the inset. **G**. Detail of two electron-dense vesicles surrounded by a double lipidic membrane (empty arrowhead) in the immediate periphery of the cell. Mitochondria (m). **H**. Unicellular stage showing detachment of the outer cell-wall. The wall presents several subtle evaginations (filled arrowhead). An electron-dense vesicle (empty arrowhead) is excreted to the space between plasma membrane and cell wall. **I**. Surface projections on a free *T. philomaios* cell (arrowheads). **J**. Parasite cell with mitochondria (m) and nucleus (n) in contact with a host cell (h). At least two flagellar structures (black arrows) have been observed flanking *T. philomaios* cells **K**. Intracellular stage of *T. philomaios* inside a host haemocyte with a thin detached wall (filled arrowhead) **L**. Coinfection of *T. philomaios* (empty arrowhead) and the ascetosporean parasite *Haplosporidium orchestiae* (filled arrowheads) in *Orchestia* sp. Only developing *T. philomaios* cells (empty arrowhead) are visible inside host haemocytes (h). Scale bars = 2 µm for (A, B, C, L), and 500 nm for (D, E, F, G, H, I, J, K). Inset in (B) is 2 µm; inset in (F) is 100 nm.

The majority of *T. philomaios* cells examined corresponded to one of the two main cell cycle stages described above. The occasional occurrence of intermediate forms and structures (Fig. 7E, 7F) suggested how unicellular cells were released from multicellular stages. The wall of the receptacle became reduced until it fractured, allowing dispersal of the walled inner cells. Just before being released, or immediately after (Fig. 7E, 7F), some of the released cells became less electron-dense, with a fine matrix between wall and plasma membrane. In later stages, the cell wall thickened and separated from the plasma membrane, possibly aided by co-occurring cellular projections (Fig. 7F). At this stage, some of the electron-dense lipid vesicles (Fig. 7G, 7H), seemingly enclosed by a double membrane, were absorbed, or excreted. Occurrence of non-walled unicellular forms of *T. philomaios* constituted the only stages in which the presence of microvilli (Fig. 7I) and maybe a flagellum (Fig. 7J) were noticeable. Inside haemocytes non-walled parasitic cells were loosely enclosed by a membrane of unknown origin (Fig. 7K). Coinfection of *T. philomaios* with *Haplosporidium* sp. (Urrutia et al. 2020) was not uncommon (Fig. 7L), but only *T. philomaios* cells were observed inside haemocytes. *In-situ* hybridization confirmed that the ultrastructure and histopathology of the amphipod infecting microeukaryotes matched with the 18S identified as *T. philomaios* (Fig. 8). The size and distribution of the DIG-NTB stained structures coincided with their immediate histological H&E stained sections. Round blue stains (2-4 µm) appeared concentrated in tegument, connective tissue, (Fig. 8A, 8B), gills, haemolymph (Fig. 8C, 8D), and inside haemocytes (Fig. 8E, 8F).

**Fig 8.**
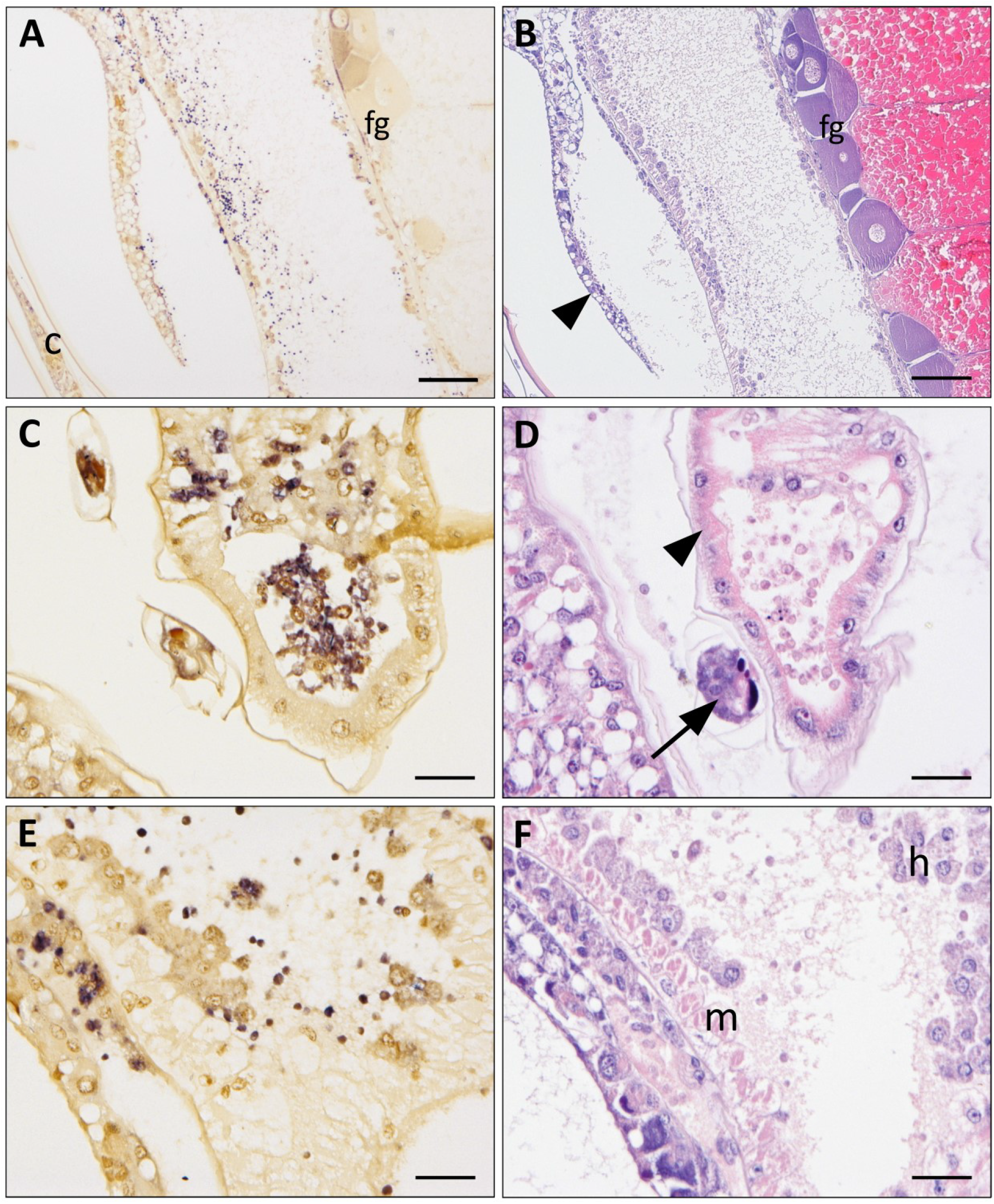
Histological sections of *Orchestia* sp. tissues following *In-Situ* Hybridization (ISH) using a DIG-labelled probe (A, C, E) and the respective consecutive histological H&E-stained section obtained from the same host (B, D, F). **A. & B**. *Txikispora philomaios* cells can be observed infecting the tegument (filled arrowhead) the cuticle (c) and haemocytes present in the cardiac tissues. Female gonads (fg) appear uninfected in this individual. **C. & D**. Infected gill cells (arrowhead) usually have ciliates attached (arrow), which are not infected in this occasion. Haemolymph circulating through the gills is heavily infected with *T. philomaios* cells. **E. & F**. Uninfected muscle (m) forming the cardiac tissue, pumps infected haemocytes (h) to other tissues. Scale bars = 100 µm for (A, B) and 25 µm for (C, D, E, F).

### Life cycle and potential vectors

The occurrence of a multicellular stage provided strong evidence that *T. philomaios* was proliferating inside amphipod hosts. Two different amphipod genera were found to be susceptible to infection by *T. philomaios*, raising questions about host specificity. Therefore, some common invertebrates cohabiting with *Echinogammarus* sp. and *Orchestia* sp. were analysed histologically and by PCR. In Newton’s Cove, co-occurring polychaetes of genus *Capitella*, the turbellarian *Procerodes* sp., and harpacticoid copepods were sampled (Table 2). While evident systemic *T. philomaios* infection in amphipods is limited to late April and May, we recognized the possibility that the parasite might be present in other hosts during a different time of the year. Thus, abundantly co-occurring invertebrates were sampled during May, June, July, August, and September. No clear evidence of *T. philomaios* cells were observed in the histopathological survey of *Procerodes* sp., *Capitella* sp. or harpacticoid copepods. However, PCR analysis carried out using sets of individuals representing these taxa indicated the presence of DNA of *T. philomaios* in a single sample comprising *Procerodes* individuals, collected during May 2019.

### Phylogenetic analyses

Initially, a partial SSU sequence (ca. 705 bp long, including variable regions V5, V7, V8, and partial V9) was coincidentally amplified by haplosporidian-specific primers (Hartikainen et al 2014) from an *Echinogammarus* sp. sample later shown to be infected by *T. philomaios*. The top Blastn match for this sequence was the ichthyosporean *Dermocystidium salmonis* (91.5% similarity; 92% coverage; e-value = 0). Phylogenetic analysis of this 705 bp sequence (not shown) placed *T. philomaios* within clade Holozoa, with low nodal support for any particular position, but often grouping with Ichthyosporea or Filasterea. A longer, equivalent 18S region of 1679 bp generated from an infected *Echinogammarus sp*. individual, resulted in a Blastn match of 87.90% similarity (99% coverage) to the free living filasterean *Pigoraptor chileana*. Phylogenetic analysis of the 1679 bp region (Fig. 9) was consistent with that using the shorter fragment, and robustly placed *T. philomaios* as an holozoan, but very weakly branching as the earliest diverging lineage in Holozoa.

**Fig 9.**
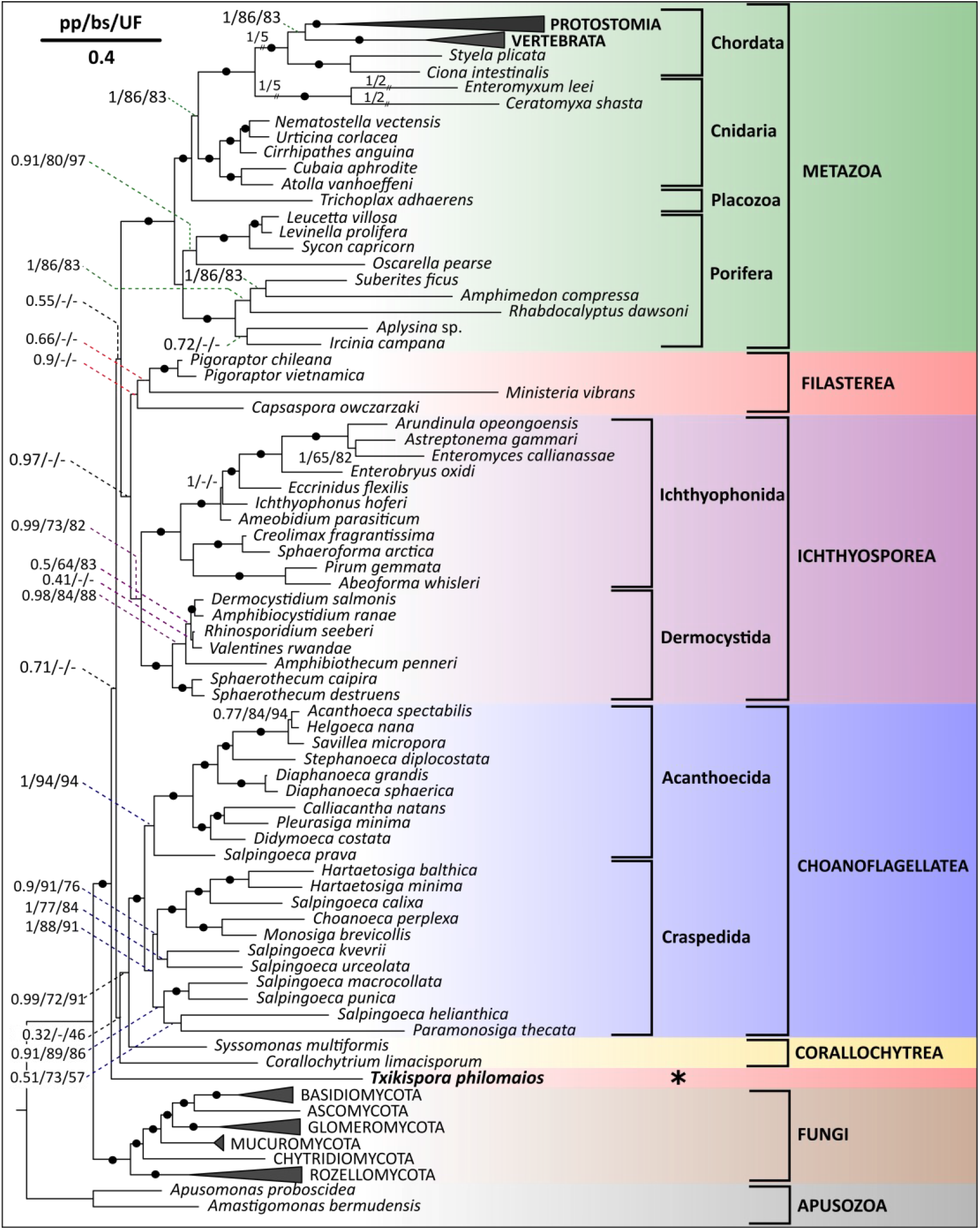
Bayesian phylogenetic analysis of 18S and 28S rRNA genes places the novel amphipod parasite *Txikispora philomaios* (1679 bp) within Holozoa. The alignment included the 1679 bp 18S rRNA gene sequence of *T. philomaios* and the 18S of the rest of species (28S sequences were included where available in Genbank). The tree includes a selection of the main opisthokont groups and unicellular holozoan lineages. Nodal support values are shown in clusters of three, representing Bayesian posterior probability (pp) run on MrBayes, maximum likelihood bootstrap support (bs) generated using RAxML with 1,000 replicates, and ML ultrafast 1,000 replicates bootstrap support (UF) from IQ-TREE, respectively. Nodes with values (> 0.95 pp, > 95% bs, > 95% UF) are represented by a black dot on the branch. Species belonging to clades Protostomia, Vertebrata, Basydiomycota, Glomeromycota, Mucuromycota, Chytridiomycota, and Rozellida were collapsed.

Several databases were mined for environmental sequences (process specified in section 2.5) related to *T. philomaios* (Table 4). The resulting phylogenetic tree (Fig. 10) showed some interesting differences when compared to the tree without environmental sequences (Fig. 9). In particular, in Fig. 10 *T. philomaios* branched within Filasterea, in a clade mostly comprising environmental sequences, but also Ministeria. The filasterean clade was more strongly supported with the inclusion of the environmental sequences, with supports of (0.98, 21, 72; posterior probability, ML bootstrap, and ML ultrafast bootstrap, respectively) compared to (0.9, -, -) in Fig. 10. The metazoan, choanoflagellate, and fungal clades were again fully/strongly supported, although the ML bootstrap support for the ichthyosporean clade was lower: 1, 34, 68 in Fig. 10 to 0.99, 73, 82 in Fig. 9. The phylogenetic position of the two pluriformean species as basal to choanoflagellates was maintained, but the support for *C. limacisporum* in that position increased from (0.32, -, 46) to (0.91, 19, 64).

**Fig 10.**
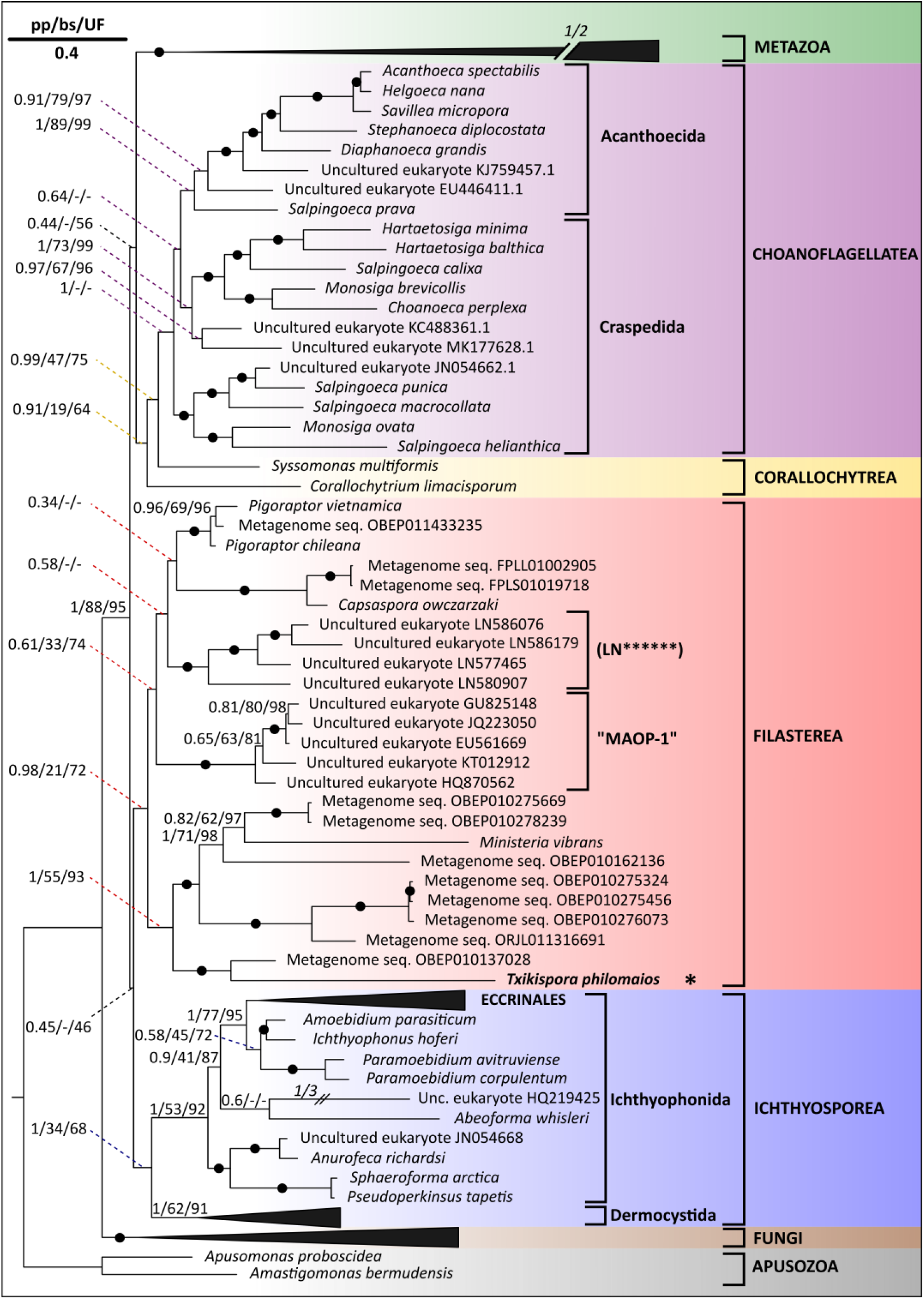
Bayesian phylogenetic analysis of 18S and 28S rRNA genes, including environmental sequences, places the novel amphipod parasite *Txikispora philomaios* (1679 bp) within Filasterea. 28S sequences were included where available in Genbank. Node support values are shown in clusters of three, representing Bayesian posterior probability (pp) run on MrBayes, maximum likelihood bootstrap support (bs) generated using RAxML, and ML ultrafast bootstrap support (UF) from IQ-TREE, respectively. Nodes with values (> 0.95 pp, > 95% bs, > 95% UF) are represented by a black dot on the branch. Species belonging to Metazoa and Fungi were collapsed, as were Eccrinales and Dermocystida (Ichthyosporea). Environmental sequences are indicated by their GenBank accession numbers.

The filasterean clade in Fig. 10 was moderately well supported by Bayesian Inference (0.98, 21, 72) but contained a large proportion of partial environmental sequences yielding disparity between ML methods. *Txikispora* was a robustly placed sister to Metagenome seq. OBEP010137028) sampled from sandy/muddy sediments associated with algae in Ulvedybet in Limfjorden (northern Denmark) (Karst et al. 2018). These, together with *Ministeria*, formed a clade with other environmental sequences from fresh groundwater systems in Denmark (OBEP010275669, OBEP010278239, OBEP010275324, OBEP010275456, OBEP010276073) and New York State (ORJL011316691) (Karst et al. 2018; Wilhelm et al. 2018), with the exception of OBEP010162136, which also came from the coastal location of Ulvedybet in Limfjorden (Karst et al. 2018). The other characterised filasterean taxa, *Capsaspora* and *Pigoraptor*, grouped separately within the filasterean clade, and potentially more closely to each other than to *Ministeria* and *Txikispora* (Fig. 10)

Several environmental sequences branched close to *Capsaspora* and *Pigoraptor*. Metagenomic sequence OBEP01433235, collected from the sediments in a freshwater lake (Denmark) was very closely related to *Pigoraptor*. Additionally, two almost identical sequences (FPLL01002905 and FPLS01019718) collected from soil samples in Denmark (Karst et al. 2016) were robustly sister to *C. owczarzaki* (100, 100, 100). Two further clades of environmental sequences branched within the filasterean clade as shown on Fig. 10. One was an abundant group of uncultured marine organisms named “MAOP1-Marine Opisthokonts”, which was weakly sister to *Pigoraptor* in Hehenberger et al. (2017). The other was a clade formed by short sequences (indicated on Fig. 10 as LN******) collected from a subterranean colony of ants adjacent to Chagres river, Panama (Scott et al. 2010).

## DISCUSSION

### Phylogeny and diversity

Until the recent addition of *Pigoraptor* by Hehenberger et al. (2017), Class Filasterea comprised only two genera: *Capsaspora* (*C. owczarzaki*) and *Ministeria* (*M. vibrans* + *M. marisola*). Hehenberger et al. (2017), also suggested the inclusion of an abundant group of marine opisthokonts “MAOP-1” (del Campo and Trillo 2013) into Filasterea. The ecology of these clade formed by uncultured organisms remains entirely undetermined except for an apparent inclination for the low oxygen fraction of the water column in coastal waters of the Indian, Atlantic and Pacific Oceans. Our results (Fig. 10) further support the inclusion of MAOP-1 in Filasterea. However, none of the ML analyses are conclusive, and the relative phylogenetic position of the group among existing filasterean species varies. In Hehenberger et al. (2017), MAOP-1 appeared as sister to *Pigoraptor* sp. (ML Bootstrap = 52%), but our analysis showed it as weakly sister to *Pigoraptor* spp, plus *C. owczarzaki* (in both cases with related environmental sequences) plus the LN****** environmental sequences. Our 18S phylogenetic analysis without environmental sequences (Fig. 9) also supported the inclusion of *T. philomaios* into Holozoa, but not its association with Filasterea. Ongoing phylogenomic analyses seek to place *T. philomaios* using a much larger number of genes.

It is well established that single-gene trees are unable to resolve deep eukaryotic phylogenetic relationships. This is particularly evident for holozoan relationships, as shown by Simion et al. (2017) among others. Our results suggest that the use of uncharacterized environmental sequences in phylogenetic studies based on 18S provide additional phylogenetic information that may assist in resolving evolutionary relationships of novel holozoan organisms, as has previously been demonstrated for other eukaryotic groups and eukaryotes as a whole (e.g. Berney et al 2004; Cavalier-Smith 2004; Bass et al. 2018; Hartikainen et al. 2016).

Several environmental sequences were closely related to existing filasterean species (Fig. 10). The uncultured sequence Metagenome seq. OBEP011433235 most likely belongs to a novel *Pigoraptor* sp. species, which evidences the preference of the genus for the sediments of stagnant freshwater systems, and a global distribution (Denmark, Chile, Vietnam). However, the environmental sequences FPLL01002905 and FPLS01019718, although sister to *C. owczarzaki*, are too distantly related to sensibly infer any lifestyle or other phenotypic similarity between them and *Capsaspora*. Interestingly, its occurrence in a Danish grassland (Karst et al. 2016), contrasts with the rest of environmental sequences associated to Filasterea, which were sampled from aquatic ecosystems. Although it is not possible to determine whether other filasterean environmental sequences are parasites, other symbionts, or free-living, our discovery of a true filasterean parasite means that this is now a realistic working hypothesis.

At some point in the evolutionary history of their lineages, *C. owczarzaki* and *T. philomaios* evolved endosymbiotic and parasitic behaviours closely associated with host haemolymph and haemocytes, highly uncommon target cells/tissues in the related clade Ichthyosporea (Glockling et al. 2013). Whether filasterean radiation preceded that of early metazoans 650-833 million years ago (Paps 2018) remains unresolved. Nonetheless, a common tissue trophism could suggest certain predisposition in the early ancestors of filastereans to colonize the haemolymph (or precursor cells) of other organisms, that could be shared by related uncultured filastereans. Actually, tissue specificity is often determined by evolutionary changes occurring early in a lineage, for instance, in ichthyosporeans a different tissue trophism allows to differentiate between the two orders (Mendoza et al. 2002), but it also happens in other prostist clades in and out Holozoa; as it is the case of myxozoans (Molnár and Székely 2014) or apicomplexans (Leander et al. 2006).

### Clinical signs and histopathology

*T. philomaios* cells congest the host’s haemolymph and tegument, making heavily infected amphipods present a light-yellow colouration and reduced carapace transparency (internal organs are not easily visible though the carapace). Definite colour alterations of the host’s carapace have been documented for other parasitic infections, such as those produced by acanthocephalans, cestodes, and trematodes (Lagrue et al. 2016; Johnson and Heard 2017). Other microeukaryotic cells targeting tegument and haemolymph in amphipods (*Haplosporidium* sp.) have also been associated with a pallid carapace and opacity. However, amphipods with heavy haplosporidiosis look whitish rather than yellowish, at least in *Echinogammarus* sp. and *Orchestia* sp. (Urrutia et al. 2019). The formation of cell aggregates, very evident in fresh haemolymph smears, are characteristic among filastereans (Sebé-Pedrós et al. 2013) and facilitates the differentiation between *T. philomaios* and other protistan parasites with similar size. We have also observed infected hosts to be more sessile and unresponsive to stimuli, but this is the case for other protist parasite infections as well, not only in amphipods (Feist et al. 2009; Lefèvre et al. 2009).

Measuring less than 3 µm in diameter *T. philomaios* is one of the smallest known holozoans. In clade Filasterea only the bacterivorous *M. vibrans* would have a similar size, with its round cells being 2.1-3.6 µm in diameter (Mylnikov et al. 2019). The highly motile predators *P. vietnamica and P. chileana* tend to be considerably bigger (5-12 µm), in the size range of most choanoflagellates and corallochytreans (Raghu-kumar 1987; Dayel and King 2014; Tikhonenkov et al. 2020). Only the zoospores of few species of ichthyosporean parasites such as *Sphaerothecum destruens* or *Dermocystidium percae* have been reported to have a similar or even smaller size than *T. philomaios* (Pekkarinen and Lotman 2003, Andreou et al. 2011). A reduced body and genome size have been linked to parasitism in other protistan groups such as microsporidians or myxozoans (Keeling and Fast 2002; Keeling 2004; Holzer et al. 2018). This has not been studied for unicellular holozoans, possibly due to the absence of parasites among choanoflagellates, and rarity of free-living forms in Ichthyosporea (Mendoza et al. 2002; Glockling et al. 2013; Hassett et al. 2015). Filasterea now includes free-living, symbiotic, and parasitic species, making it a good candidate for such comparative analyses, especially when ecological and genomic data from the group’s uncharacterised diversity are elucidated. The presence of holozoan protists with reduced genomes could provide very valuable information, as many studies focus on gene gains and losses to understand how and when animal multicellularity evolved (Paps et al. 2013; Grau-Bove et al. 2017; Richter et al. 2018).

### Ultrastructure

Ultrastructurally, a crown of microvilli around a single flagellum makes choanoflagellates the most easily identifiable of all unicellular holozoans (Mah et al. 2014). The presence of a single posterior flagellum is a hallmark trait among opisthokonts (Cavalier-Smith 1987) and also been observed in holozoan Classes Filasterea, Corallochytrea and Ichthyosporea (Marshall et al. 2008; Torruella et al. 2015; Hehenberger et al. 2017; Mylnikov et al. 2019).

Fresh smears of *T. philomaios* showed the presence of cell-projections comparable to the flagellar structures described by light microscopy and TEM in *M. vibrans* and *Pigoraptor* sp. (Torruella et al. 2015; Hehenberger et al. 2017; Mylnikov et al. 2019). However, no evidence of a flagellum was observed in the histopathological analysis, and we only have limited ultrastructural evidence of its occurrence by TEM (Fig. 7J). While inconclusive, we must note that in fresh smears *T. philomaios* cells were exposed to a substrate and marine water, but histology and TEM analysed them fixed in tissues and haemolymph. The zoospores of dermocystids are the only known flagellated stage among parasitic/endosymbiotic holozoan protists, and quickly lose the flagellum after penetrating into the host (Pekkarinen et al. 2003). Besides, the flagellum of *M. vibrans* was only observed after examination of over 1,000 cells (Mylnikov et al. 2019), a number not reached for *T. philomaios*, which has also resisted culturing attempts (see below). A non-flagellated *T. philomaios* would imply a secondary loss of the structure (based in our phylogeny, Fig. 10), the second one within Filasterea after *C. owczarzaki*. Two losses are less parsimonious, but could strengthen the idea of a parasitic/endosymbiotic lifestyle driving them, which has also been suggested for non-flagellated ichthyosporean parasites in order Ichthyophonida (Marshall and Berbee 2011, Torruella et al. 2015).

Microvilli are actin-based filopodial structures present in filozoans (Karpov et al. 2016; Sebé-Pedrós et al. 2017; Mylnikov et al. 2019). They form a crown around the flagellum in choanoflagellates they are evenly distributed around the cell in all filastereans and pluriformeans (Mylnikov et al. 2019; Tikhonenkov et al. 2020), clades in which they can be up to three or four times the length of the cell (10 µm in *M. vibrans*, 20 µm in *C. owczarzaki*, and 34 µm in *S. multiformis*). However, they are not present in cystic and dividing stages, what could explain the reduced evidence for them in *T. philomaios* (Fig. 7I). Moreover, their occurrence was not noticed in the original descriptions of *C. owczarzaki* done on explanted pericardial sacs of snails (Owczarzak et al. 1980), but they are evident when the facultative symbiont is in axenic culture (Sebé-Pedrós et al. 2013), where they have been shown to facilitate movement, cell-cell adhesion, and food particle capture (Parra-Acero et al. 2020). It is possible that microvilli are not desirable in the haemolymph of a host, where the current impedes movement and there is no substrate surface other than target haemolymph cells.

Opisthokonts are characterized by flat non-discoid cristae (Cavalier-Smith and Chao 1995), with ichthyosporean *Ichthyophonus hoferi* being one of the few exceptions (Spanggaard and Huss 1996). Mitochondria in *T. philomaios* follows the norm and possesses lamellar cristae (Fig. 7G, 7I). The radial distribution of numerous mitochondria in the periphery of non-cystic stages (Fig. 6A) could indicate a close in time cell division between daughter cells, as observed in the ichthyosporean *Sphaerothecum* sp. (Borteiro et al. 2018). In contrast, the absence of mitochondria in stages with a thicker wall suggests a resistant spore-like stage, as it is the case in the ichthyosporean *Amphibiocystidium* sp. (González-Hernández et al. 2010). However, the structure and activity of mitochondria in parasites has been observed to be extremely flexible (Zíková et al. 2016), as they would be able to use mitochondrial metabolites of the host (de Melo and Souza 1992).

Numerous electron-dense bodies comparable to those observed in other filasterean species (Owczarzak et al. 1980, Tikhonenkov et al. 2020) are scattered in the cytoplasm of *T. philomaios* (Fig. 6A, 6C, 6D, 7G, 7H, 7K). However, their occurrence is not characteristic of filastereans or even holozoan protists, as they have been observed in distantly related clades such as apicomplexans, ascetosporeans or dinoflagellates (Speer et al. 1999; Stentiford and Shields 2005; Feist et al. 2009). Nevertheless, their size and number has been suggested to be of taxonomic value in Mesomycetozoea (Pereira et al. 2005), and indicative of the function of certain life stages and their maturation (Vilela and Mendoza 2012; Fagotti et al. 2020). These bodies have been described as lipid globules in *M. vibrans* (Mylnikov et al. 2019) and reserve substances (most likely glycoprotein) in genera *Pigoraptor* and *Syssomonas* (Tikhonenkov et al. 2020). In contrast, the occurrence of a double lipidic layer around them in *C. owczarzaki* made Owczarzak et al. (1980) suggest that these “lipid filled vacuoles” were excreted. In *T. philomaios* we observe two main forms; the first is a smaller and electron-lucent body similar to those observed in genera *Ministeria, Pigoraptor*, and *Syssomonas*. The second form is a larger and electron-dense body surrounded by a double lipid layer (Fig. 6H, 7G) that appears to be excreted (Fig. 7H) as proposed for *C. owczarzaki*. However, its implication in the formation of the cell wall should be considered, as it is not clear how the ejected material could trespass the outer membrane (Fig. 7G).

### Life cycle and potential hosts

So far, all filastereans have been culturable (Stibbs et al. 1979; Cavalier-Smith and Chao 2003; Hehenberger et al. 2017; Mylnikov et al. 2019), allowing a detailed description of their life cycle in culture conditions. In contrast, *T. philomaios*, like most parasites in the clade Ichthyosporea remains unculturable (Cafaro 2005; Glockling et al. 2013). According to the diagnostic description of Class Filasterea Cavalier-Smith 2008, trophic stages in this lineage do not possess a cell wall (Shalchian-Tabrizi et al. 2008). In free-living genera *Pigoraptor* and *Ministeria*, this non-walled stage corresponds to a flagellated amoeba which uses its retractile microvilli to capture preys and attract food particles (Hehenberger et al. 2017; Mylnikov et al. 2019). In turn, trophocytes of the endosymbiont *C. owczarzaki* lack a flagellum, and even microvilli if cultured in explanted tissues of *B. glabrata* (Owczarzak et al. 1980; Sebé-Pedrós et al. 2013). Although morphologically different, the behaviour of trophocytes is the same in all known filasterean species; they can either divide by binary fission or encyst when the food source is depleted (Hertel et al. 2002; Tikhonenkov et al. 2020). The binary fission observed by light microscopy in few walled cells of *P. vietnamica* (Tikhonenkov et al. 2020) represents the only known exception of cellular division occurring outside the trophic stage. Interestingly, our TEM analysis indicates that quite the contrary occurs for *T. philomaios*, in which cell division appears to occur exclusively inside walled cells (Fig. 6B, 6I, 7D) as in Corallochytrea and Ichcthyosporea (Raghu-Kumar 1987; Lotman et al. 2000; Pekkarinen et al. 2003; Glockling et al. 2013). If flagella and/or microvilli occur in *T. philomaios* trophocytes (Fig. 7I, 7J), these structures are lost when parasitic cells either penetrate or are engulfed by host haemocytes (Fig. 7K).

A single host haemocyte can contain up to ten *T. philomaios* cells, in which four walled endospores arise inside walled parent cells (Fig. 7D). Comparable cellular structures containing 16-32 endospores are the result of a palintomic division in corallochytrean cystic stages (Raghu-Kumar et al. 1987; Tikhonenkov et al. 2020). Once mature, *T. philomaios* endospores would leave the parent cells through an opening formed in its wall, by which time its thickness is much reduced, as in Corallochytrea and Ichthyosporea (Mendoza et al. 2002; Marshall and Berbee 2011; Tikhonenkov et al. 2020). The wall thickness, electron-density and amount of reserve material vary greatly among endospores. Some cells appear active even before exiting the ruptured parent cell (Fig. 7E), presumably ready to re-infect other haemocytes and tissues in the same host, as it has been shown for several ichthyosporeans (Arkush et al. 2003; Marshall et al. 2008; Kocan 2019). Other cysts seem to be resistant (Fig. 6D, 6F), perhaps capable of infecting other amphipods or even remaining viable in the environment for months (Marshall and Berbee 2010; Gozlan et al. 2014; LaPatra and Kocan 2016).

The transmission method for *T. philomaios* cells is unknown, as for *C. owczarzaki* (Harcet et al. 2016), and most parasites in Ichthyosporea (Glockling et al. 2013). A direct cycle by consumption of infected prey has been demonstrated in the ichthyophonid *I. hofferi* (Kramer-Schadt et al. 2010), and could be possible for *T. philomaios*, given the high levels of interspecific predation (Dick et al. 1999), cannibalism (Kinzler and Maier 2003), and scavenging of conspecifics (Agnew and Moore 1986) observed in amphipods. The thicker ameboid endospores observed in *T. philomaios* are also remindful of the infective waterborne cells observed in ichthyophonid parasites (Olson et al. 1991, Andreou et al. 2009, Kocan 2019), which unlike those in order Dermocystida, lack a flagellum (Mendoza et al. 2002). Additionally, cysts of the so called “™S” ichthyosporean infecting *Tenebrio molitor*, persist in the connective tissues associated to the gonads, and are transmitted with sperm during copulation (Lord et al. 2012). The presence of few *T. philomaios* cells infecting amphipod gonads throughout the year (although with low prevalence = 1.9%) leaves open the possibility of a similar “nuptial transmission” for the novel parasite. In that case, *Echinogammarus* sp. and *Orchestia* sp. would represent the reservoir for *T. philomaios*, which according to the most extended definition is an environment/population where the pathogen can be permanently maintained and transmitted (Haydon et al. 2002).

Finally, an indirect transmission cycle has been contemplated as well, given the generalist infectivity observed in *T. philomaios* and ichthyosporean parasites (Andreou et al. 2012; Rowley et al. 2013; Combe and Gozlan 2018). Copepods have been proposed as the missing intermediate host for the fish parasite *I. hofferi*, which infects herring and salmon species (Hershberger et al. 2002; Gregg et al. 2012). Interestingly, harpacticoid copepods are some of the most common invertebrates co-occurring with amphipods in the upper part of the intertidal in Newton’s Cove, Camel, Dart and Tamar estuaries (personal observation; Hicks and Coull 1983). However, our PCR based search for *T. philomaios* in copepods (n = 1300 individuals) was negative, just like the histopathological analysis. In turn, the results for the turbellarian *Procerodes* sp. were PCR positive during May. The platyhelminth, which is very common in the north Atlantic, appears to predate on diseased *Echinogammarus* sp. preys and carcasses (Den Hartog 1968; Taylor 1986), showing a link and a possible role as intermediate host. A more extensive histopathological analysis of *Procerodes* sp. will be necessary to substantiate its possible role as intermediate host of *T. philomaios*. If uninfected the turbellarian could still be a vector helping the dispersal of viable *T. philomaios* cysts.

### Distribution, prevalence, and ecological significance

The low number of filasterean species and their rare appearance in environmental samplings have prevented any previous estimation of their temporal prevalence, as it has been assayed for larger holozoan clades Ichthyosporea and Choanoflagellatea (Marchant and Perrin 1990; Kasesalu et al. 2000; Pekkarinen and Lotman 2003). The prevalence of *C. owczarzaki* in *Biomphalaria glabrata* has been observed to vary from 1% to 45% depending on the strain (Hertel et al. 2002), but the measurement, done on cultured snails, does not estimate occurrence on a time period. Our study is the first one to reveal a temporal pattern in the abundance of a filasterean species. The quickly vanishing peak in prevalence observed for *T. philomaios* during May, exposes seasonality as an until now unaccounted bias for the scarcity of filasterean sequences in environmental samplings (del Campo et al. 2015; Hehenberger et al. 2017; Mylnikov et al. 2019). A similar short temporary window in the transmission of *C. owczarzaki* between snails, could explain, at least partially how it has eluded sampling efforts to find it in the wild (Ferrer-Bonet and Ruiz-Trillo 2017). Additionally, our failed efforts to amplify the 18S of *T. philomaios* from filtered water collected in Newton’s Cove during May, reinforces the hypothesis of a reduced detection capability of eDNA for parasites/endosymbionts (Dumonteil et al. 2018).

So far, it has been observed that *T. philomaios* is able to infect at least two different amphipod genera, indicating certain range of hosts specificity that could expand notably if infection in the turbellarian *Procerodes* sp. is substantiated by histology. In this study, the prevalence of *T. philomaios* was as high as 64% (May 2016), with about a third of the infected individuals presenting heavy infections associated to tissue disruption and haemolymph congestion by parasitic cells. From the point of view of pathology, other protistan parasites that tend to multiply and congest the haemolymph of crustacean hosts, such as the dinoflagellate *Hematodinium* sp. have been associated with a reduced oxygenation capability and diminished overall fitness (Taylor et al. 1996; Stentiford et al. 2001). The observed unresponsiveness to stimuli in infected amphipods, is consistent with the systemic damage observed in the tegument, which functions as the sensorial system (Steele and Oshel 1987). Collectively, numerous protistan parasites have been found to profoundly alter the populations of amphipods and other crustaceans (Morado 2011; Ironside and Alexander 2015). Considering that several ichthyosporean parasites are responsible for important mortality events in fish and amphibian populations (Raffel et al. 2008; Kirkbright et al. 2016) it would be interesting to monitor the influence of *T. philomaios* in the amphipod population, as *Echinogammarus* and *Orchestia* are amongst the most common and abundant crustaceans in coastal ecosystems of Northern Europe (Marques and Nogueira 1991; Mantzouki et al. 2012), and important invasive species outside the continent (Van Overdijk et al. 2003; Herkül et al. 2006).

## TAXONOMIC SUMMARY

Eukaryota Chatton, 1925 / Eukarya Margulis and Chapman, 2009: Opisthokonta Adl, 2005: Holozoa Adl, 2012: Filasterea Shalchian-Tabrizi, 2008: Ministerida Cavalier-Smith, 1997

### Family Txikisporidae Urrutia, Feist & Bass n. fam

*Diagnosis*. Naked unicellular and uninucleated protists morphologically similar to individuals in family Ministeriidae Cavalier-Smith 2008, but with a parasitic lifestyle.

*Type genus. Txikispora* n. g. (see below)

### Genus *Txikispora* Urrutia, Feist&Bass n. g

*Etymology*. ‘txiki’: small and ‘spora’: a seed (Basque). The name has been chosen to reflect relatedness with the filasterean endosymbiont *Capsaspora* Hertel, 2002 (“the quick eating seed”) and its small size, while putting a distance with other small spore forming parasitic lineages with Latin stems.

*Diagnosis*. As for species (see below)

*Type species. Txikispora philomaios* (see below)

### *Txikispora philomaios* Urrutia, Feist &Bass n. sp

*Etymology*. Txiki-: small, spora: spore, philo-: lover, maios: the month of May. “The little May-loving spore”, referring to its predominant detection (as a parasite of amphipods) in that month. *Diagnosis*. Virtually spherical monokaryotic stages, with a length of 2.6 ± 0.41 µm and a width of 2.26 ± 0.34 µm. The round and walled multinucleated stage contains four walled cells inside, which resemble a lot the monokaryotic stages. The size of this divisional stage is slightly bigger (3.17 µm ± 0.24 in diameter). Infection develops principally inside host haemocytes and connective tissues, especially those associated to the tegument. Infection in amphipods in the southwest of UK occurs consistently during late April and May, the prevalence of the parasite during the rest of the year is anecdotical (< 2%). The parasite has been also linked to the gonads, being the only organ that appears to be infected during the rest of the year. There is host reaction to the parasite in form of melanization and granuloma formation, especially when the parasite affects the hepatopancreas.

*Type host*. Amphipods *Echinogammarus* sp. and *Orchestia* sp.

*Type location*. Coastal waters in Newton’s Cove (UK)

*Type material*. Original slides used for this paper are stored together with biological material embedded in wax and epoxy resin in Cefas Weymouth Lab. Type material is stored as RA16020 (specimen no. 19) and RA17028 (specimen no. 53) and (specimen no. 287). The SSU rDNA sequence is deposited in GenBank under accession number (to be submitted).

## ACKNOWLEDGMENTS

A.U was supported by a PhD studentship grant (Programa Predoctoral de Formación de Personal Investigador – Departamento de Educación Gobierno Vasco) in collaboration with Ikerbasque; the experimental work was funded by the UK Department of Environment, Food and Rural Affairs (Defra) under contracts FC1214 (to D.B) and FB002A (to S.W.F and D.B). IR-T was supported by a grant (BFU2017-90114-P) from Ministerio de Economía y Competitividad (MINECO), Agencia Estatal de Investigación (AEI), and Fondo Europeo de Desarrollo Regional (FEDER). M.M.L. was supported by a Marie Skłodowska-Curie Individual Fellowship under the EU Framework Programme for Research and Innovation Horizon 2020 (Project ID 747789), and an Ayuda Juan de la Cierva -Incorporación (IJC2018-036657-I).We thank Matt Green, Rose Kerr, Dr. Corey Holt, Dr. Georgia Ward, Dr. Selina Church, and Caroline Daumich, who assisted with sample collection and processing and provided advise.

## LITERATURE CITED

1. Agnew, D. J. & Moore, P. G. 1986. The feeding ecology of two littoral amphipods (Crustacea), Echinogammarus pirloti (Sexton & Spooner) and E. obtusatus (Dahl). J. Exp. Mar. Biol. Ecol., 103:203–215.

2. Amaral-Zettler, L. A., Nerad, T. A., O’Kelly, C. J. & Sogin, M. L. 2001. The nucleariid amoebae: more protists at the animal-fungal boundary. J. Eukaryotic Microbiol., 48:293–297.

3. Andreou, D., Arkush, K. D., Guégan, J. F. & Gozlan, R. E. 2012. Introduced pathogens and native freshwater biodiversity: a case study of Sphaerothecum destruens. PLoS One., 7, e36998. doi:10.1371/journal.pone.0036998

4. Andreou, D., Gozlan, R. E. & Paley, R. 2009. Temperature influence on production and longevity of Sphaerothecum destruens zoospores. J. Parasitol., 95:1539–1541.

5. Andreou, D., Gozlan, R. E., Stone, D., Martin, P., Bateman, K. & Feist, S. W. 2011. Sphaerothecum destruens pathology in cyprinids. Dis. Aquat. Org., 95:145–151.

6. Arkush, K. D., Mendoza, L., Adkison, M. A. & Hedrick, R. P. 2003. Observations on the life stages of Sphaerothecum destruens n. g., n. sp., a mesomycetozoean fish pathogen formally referred to as the rosette agent. J. Eukaryotic Microbiol., 50:430–438.

7. Arroyo, A. S., López-Escardó, D., Kim, E., Ruiz-Trillo, I. & Najle, S. R. 2018. Novel diversity of deeply branching Holomycota and unicellular holozoans revealed by metabarcoding in Middle Paraná River, Argentina. Front. Ecol. Evol., 6:99.

8. Bancroft, J. D. & Cook, H. C. 1994. Manual of Histological Techniques and their Diagnostic Applications, Churchill Livingstone, New York, NY. p. 40–41.

9. Bass D., Tikhonenkov, D. V., Foster, R., Dyal, P., Janouskovec, J., Keeling, P. J., Gardner, M., Neuhauser, S., Hartikainen, H., Mylnikov, A. P. & Berney, C. 2018. Rhizarian ‘Novel Clade 10’ revealed as abundant and diverse planktonic and terrestrial flagellates, including Aquavolon n. gen. J. Eukaryot. Microbiol., 65:828–842.

10. Bass, D., Yabuki, A., Santini, S., Romac, S. & Berney, C. 2012. Reticulamoeba is a long-branched Granofilosean (Cercozoa) that is missing from sequence databases. PLoS One, 7, e49090. doi:10.1371/journal.pone.0049090

11. Berney, C., Fahrni, J. & Pawlowski, J. 2004. How many novel eukaryotic ‘kingdoms’? Pitfalls and limitations of environmental DNA surveys. BMC Biol., 2:13.

12. Borteiro, C., Baldo, D., Maronna, M. M., Baeta, D., Sabbag, A. F., Kolenc, F., Debat, C. M., Haddad, C. F. B., Cruz, J. C., Verdes, J. M. & Ubilla, M. 2018. Amphibian parasites of the Order Dermocystida (Ichthyosporea): current knowledge, taxonomic review and new records from Brazil. Zootaxa, 4461:499–518.

13. Broz, O. & Privora, M. 1952. Two skin parasites of Rana temporaria: Dermocystidium ranae Guyénot & Naville and Dermosporidium granulosum n. sp. Parasitology, 42:65–69.

14. Cafaro, M. J. 2005. Eccrinales (Trichomycetes) are not fungi, but a clade of protists at the early divergence of animals and fungi. Mol. Phylogenet. Evol., 35:21–34.

15. Capella-Gutiérrez, S., Silla-Martínez, J. M. & Gabaldón, T. 2009. trimAl: a tool for automated alignment trimming in large-scale phylogenetic analyses. Bioinformatics, 25:1972–1973.

16. Cavalier-Smith, T. 1987. The simultaneous symbiotic origin of mitochondria, chloroplasts, and microbodies. Ann. N. Y. Acad. Sci., 503:55–71.

17. Cavalier-Smith, T. 1993. Kingdom protozoa and its 18 phyla. Microbiol. Mol. Biol. Rev., 57:953–994.

18. Cavalier-Smith, T. & Chao, E. E. 1995. The opalozoan Apusomonas is related to the common ancestor of animals, fungi, and choanoflagellates. Proc. R. Soc. London, Ser. B, 261:1–6.

19. Cavalier-Smith, T. & Chao, E. E. Y. 2003. Phylogeny of choanozoa, apusozoa, and other protozoa and early eukaryote megaevolution. J. Mol. Evol., 56:540–563.

20. Cavalier-Smith, T. 2004. Only six kingdoms of life. Proc. R. Soc. London, Ser. B, 271:1251–1262.

21. Colley, D. G., Bustinduy, A. L., Secor, W. E. & King, C. H. 2014. Human schistosomiasis. Lancet, 383:2253–2264.

22. Combe, M. & Gozlan, R. E. 2018. The rise of the rosette agent in Europe: An epidemiological enigma. Transboundary Emerging Dis., 65:1474–1481.

23. Dayel, M. J. & King, N. 2014. Prey capture and phagocytosis in the choanoflagellate Salpingoeca rosetta. PLoS One, 9, e95577. doi:10.1371/journal.pone.0095577

24. de Melo, E. J. T. & de Souza, W. 1992. Penetration of Toxoplasma gondii into host cells induces changes in the distribution of the mitochondria and the endoplasmic reticulum. Cell Struct. Funct., 17:311–317.

25. del Campo, J. & Ruiz-Trillo, I. 2013. Environmental survey meta-analysis reveals hidden diversity among unicellular opisthokonts. Mol. Biol. Evol., 30:802–805.

26. del Campo, J., Mallo, D., Massana, R., de Vargas, C., Richards, T. A. & Ruiz-Trillo, I. 2015. Diversity and distribution of unicellular opisthokonts along the European coast analysed using high-throughput sequencing. Environ. Microbiol., 17:3195–3207.

27. Den Hartog, C. 1968. Marine triclads from the Plymouth area. J. Mar. Biol. Assoc. U. K., 48:209–223.

28. Dick, J. T., Montgomery, W. I. & Elwood, R. W. 1999. Intraguild predation may explain an amphipod replacement: evidence from laboratory populations. J. Zool., 249:463–468.

29. Dumonteil, E., Ramirez-Sierra, M. J., Pérez-Carrillo, S., Teh-Poot, C., Herrera, C., Gourbière, S. & Waleckx, E. 2018. Detailed ecological associations of triatomines revealed by metabarcoding and next-generation sequencing: implications for triatomine behavior and Trypanosoma cruzi transmission cycles. Sci. Rep., 8:1–13.

30. Eveland, L. K. & Haseeb, M. A. 2011. Laboratory rearing of Biomphalaria glabrata snails and maintenance of larval schistosomes in vivo and in vitro. In: Toledo, R. & Fried, B. (ed.), Biomphalaria snails and larval trematodes. Springer, New York, NY. p. 33-35

31. Fagotti, A., Rossi, R., Paracucchi, R., Lucentini, L., Simoncelli, F. & Di Rosa, I. 2020. Developmental stages of Amphibiocystidium sp., a parasite from the Italian stream frog (Rana italica). Zoology (Jena), 141, e125813. doi:10.1016/j.zool.2020.125813

32. Feist, S. W., Hine, P. M., Bateman, K. S., Stentiford, G. D. & Longshaw, M. 2009. Paramarteilia canceri sp. n. (Cercozoa) in the European edible crab (Cancer pagurus) with a proposal for the revision of the order Paramyxida Chatton, 1911. Folia Parasitol., 56:73.

33. Ferrer-Bonet, M. & Ruiz-Trillo, I. 2017. Capsaspora owczarzaki. Curr. Biol., 27:829–830.

34. Fredricks, D. N., Jolley, J. A., Lepp, P. W., Kosek, J. C. & Relman, D. A. 2000. Rhinosporidium seeberi: a human pathogen from a novel group of aquatic protistan parasites. Emerging Infect. Dis., 6:273

35. Glockling, S. L., Marshall, W. L. & Gleason, F. H. 2013. Phylogenetic interpretations and ecological potentials of the Mesomycetozoea (Ichthyosporea). Fungal Ecol., 6:237–247.

36. González-Hernández, M., Denoël, M., Duffus, A. J., Garner, T. W., Cunningham, A. A. & Acevedo-Whitehouse, K. 2010. Dermocystid infection and associated skin lesions in free-living palmate newts (Lissotriton helveticus) from Southern France. Parasitol. Int., 59:344–350.

37. Gouy, M., Guindon, S. & Gascuel, O. 2010. SeaView version 4: a multiplatform graphical user interface for sequence alignment and phylogenetic tree building. Mol. Biol. Evol., 27:221–224.

38. Gozlan, R. E., Marshall, W., Lilje, O., Jessop, C., Gleason, F. H. & Andreou, D. 2014. Current ecological understanding of fungal-like pathogens of fish: what lies beneath? Front. Microbiol., 5:62.

39. Grau-Bove, X., Torruella, G., Donachie, S., Suga, H., Leonard, G., Richards, T. A. & Ruiz-Trillo, I. 2017. Dynamics of genomic innovation in the unicellular ancestry of animals. eLife, 6, e26036. doi:10.7554/eLife.26036.

40. Gregg J., Grady C., Friedman C. & Hershberger P. 2012. Inability to demonstrate fish-to-fish transmission of Ichthyophonus from laboratory infected Pacific Herring Clupea pallasii to naïve conspecifics. Dis. Aquat. Org., 99:139–144.

41. Harcet, M., Lopez-Escardo, D., Sebe-Pedros, A. & Ruiz-Trillo, I. 2016. Predatory capabilities of the filasterean Capsaspora owczarzaki reveals its potential for a free-living lifestyle. Protistology, 10:26

42. Hartikainen, H., Bass, D., Briscoe, A. G., Knipe, H., Green, A. J. & Okamura, B. 2016. Assessing myxozoan presence and diversity using environmental DNA. Int. J. Parasitol., 46:781–792.

43. Hassett, B. T., López, J. A. & Gradinger, R. 2015. Two new species of marine saprotrophic sphaeroformids in the Mesomycetozoea isolated from the sub-Arctic Bering Sea. Protist, 166:310–322.

44. Haydon, D. T., Cleaveland, S., Taylor, L. H. & Laurenson, M. K. 2002. Identifying reservoirs of infection: a conceptual and practical challenge. Emerging Infect. Dis., 8:1468–1473.

45. Heger, T. J., Giesbrecht, I. J., Gustavsen, J., del Campo, J., Kellogg, C. T., Hoffman, K. M., Lertzman, K., Mohn, W. W. & Keeling, P. J. 2018. High-throughput environmental sequencing reveals high diversity of litter and moss associated protist communities along a gradient of drainage and tree productivity. Environ. Microbiol., 20:1185–1203.

46. Hehenberger, E., Tikhonenkov, D. V., Kolisko, M., Del Campo, J., Esaulov, A. S., Mylnikov, A. P. & Keeling, P. J. 2017. Novel predators reshape holozoan phylogeny and reveal the presence of a two-component signaling system in the ancestor of animals. Curr. Biol., 27:2043–2050.

47. Herkül, K., Kotta, J. & Kotta, I. 2006. Distribution and population characteristics of the alien talitrid amphipod Orchestia cavimana in relation to environmental conditions in the Northeastern Baltic Sea. Helgol. Mar. Res., 60:121.

48. Hershberger, P. K., Stick, K., Bui, B., Carroll, C., Fall, B., Mork, C., Perry, J. A., Sweeney, E., Wittouck Winton, J. & Kocan, R. 2002. Incidence of Ichthyophonus hoferi in Puget Sound fishes and its increase with age of Pacific herring. J. Aquat. Anim. Health, 14:50–56.

49. Hertel, L. A., Barbosa, C. S., Santos, R. A. L. & Loker, E. S. 2004. Molecular identification of symbionts from the pulmonate snail Biomphalaria glabrata in Brazil. J. Parasitol., 90:759–763.

50. Hertel, L. A., Bayne, C. J. & Loker, E. S. 2002. The symbiont Capsaspora owczarzaki, nov. gen. nov. sp., isolated from three strains of the pulmonate snail Biomphalaria glabrata is related to members of the Mesomycetozoea. Int. J. Parasitol., 32:1183–1191.

51. Hibberd, D. 1975. Observations on the ultrastructure of the choanoflagellate Codosiga botrytis Saville-Kent with special reference to the flagellar apparatus. J. Cell Sci., 17:191–219.

52. Hicks, G. F. & Coull, B. C. 1983. The ecology of marine meiobenthic harpacticoid copepods. Oceanogr. Mar. Biol., 21:67–175.

53. Holzer, A. S., Bartošová-Sojková, P., Born-Torrijos, A., Lövy, A., Hartigan, A. & Fiala, I. 2018. The joint evolution of the Myxozoa and their alternate hosts: a cnidarian recipe for success and vast biodiversity. Mol. Ecol., 27:1651–1666.

54. Hopwood, D. 1969. Fixatives and fixation: a review. Histochem J., 1:323–360.

55. Ironside, J. E. & Alexander, J. 2015. Microsporidian parasites feminise hosts without paramyxean co-infection: support for convergent evolution of parasitic feminisation. Int. J. Parasit., 45:427–433.

56. James-Clark, H. 1868. On the Spongiae ciliatae as Infusoria flagellata; Or observations on the structure, animality, and relationship of Leucosolenia botryoides, Bowerbank. Ann. Mag. Nat. Hist., 1:188–215.

57. Johnson, D. S. & Heard, R. 2017. Bottom-up control of parasites. Ecosphere, 8, e01885. doi:10.1002/ecs2.1885

58. Kalyaanamoorthy, S., Minh, B. Q., Wong, T. K., von Haeseler, A. & Jermiin, L. S. 2017. ModelFinder: fast model selection for accurate phylogenetic estimates. Nat. methods, 14:587–589.

59. Karpov, S., Mamkaeva, M. A., Aleoshin, V., Nassonova, E., Lilje, O. & Gleason, F. H. 2014. Morphology, phylogeny, and ecology of the aphelids (Aphelidea, Opisthokonta) and proposal for the new superphylum Opisthosporidia. Front. Microbiol., 5:112.

60. Karst, S. M., Dueholm, M. S., McIlroy, S. J., Kirkegaard, R. H., Nielsen, P. H. & Albertsen, M. 2018. Retrieval of a million high-quality, full-length microbial 16S and 18S rRNA gene sequences without primer bias. Nat. Biotechnol., 36:190.

61. Karst, S. M., Dueholm, M. S., McIlroy, S. J., Kirkegaard, R. H., Nielsen, P. H. & Albertsen, M. 2016. Thousands of primer-free, high-quality, full-length SSU rRNA sequences from all domains of life. BioRxiv, 070771. doi:10.1101/070771

62. Kasesalu, J., Laius, A. & Lotman, K. 2000. The occurrence and species composition of parasites of the genus Dermocystidium in Estonian fish farms and some natural water bodies. Agraarteadus, 11:205–212.

63. Katoh, K., Rozewicki, J. & Yamada, K. D. 2019. MAFFT online service: multiple sequence alignment, interactive sequence choice and visualization. Briefings Bioinf., 20:1160–1166.

64. Keeling, P. J. 2004. Reduction and compaction in the genome of the apicomplexan parasite Cryptosporidium parvum. Dev. Cell, 6:614–616.

65. Keeling, P. J. & Fast, N. M. 2002. Microsporidia: biology and evolution of highly reduced intracellular parasites. Annu Rev. Microbiol., 56:93–116.

66. King, N. 2004. The unicellular ancestry of animal development. Dev. Cell, 7:313–325.

67. King, N. 2005. Choanoflagellates. Curr. Biol., 15:113–114.

68. Kinzler, W. & Maier, G. 2003. Asymmetry in mutual predation: possible reason for the replacement of native gammarids by invasives. Arch. Hydrobiol., 157:473–481.

69. Kirkbright, D., Huber, P., Lillie, B. N. & Lumsden, J. S. 2016. Dermocystidium-like organism linked with a mortality event in Yellow Perch Perca flavescens (Mitchill) in Ontario, Canada. J. Fish Dis., 39:597–601.

70. Kocan, R. M. 2019. Transmission models for the fish pathogen Ichthyophonus: synthesis of field observations and empirical studies. Can. J. Fish. Aquat. Sci., 76:636–642.

71. Kramer-Schadt, S., Holst, J. C. & Skagen, D. 2010. Analysis of variables associated with the Ichthyophonus hoferi epizootics in Norwegian spring spawning herring, 1992–2008. Can. J. Fish. Aquat. Sci., 67:1862–1873.

72. Lagrue, C., Heaphy, K., Presswell, B. & Poulin, R. 2016. Strong association between parasitism and phenotypic variation in a supralittoral amphipod. Mar. Ecol.: Prog. Ser., 553:111–123.

73. LaPatra, S. E. & Kocan, R. M. 2016. Infected donor biomass and active feeding increase waterborne transmission of Ichthyophonus sp. to Rainbow trout sentinels. J. Aquat. Anim. Health, 28:107–113.

74. Laval, M. 1971. Ultrastructure et mode de nutrition du choanoflagellé Salpingoeca pelagica, sp. nov. Comparaison avec les choanocytes des spongiaires. Protistologica, 7:325–336.

75. Leander, B. S., Lloyd, S. A., Marshall, W. & Landers, S. C. 2006. Phylogeny of marine gregarines (Apicomplexa) - Pterospora, Lithocystis and Lankesteria - and the origin(s) of coelomic parasitism. Protist, 157:45–60.

76. Lefèvre, T., Lebarbenchon, C., Gauthier-Clerc, M., Misse, D., Poulin, R. & Thomas, F. 2009. The ecological significance of manipulative parasites. Trends Ecol. Evol., 24:41–48.

77. López-Escardó, D., Grau-Bové, X., Guillaumet-Adkins, A., Gut, M., Sieracki, M. E. & Ruiz-Trillo, I. 2019. Reconstruction of protein domain evolution using single-cell amplified genomes of uncultured choanoflagellates sheds light on the origin of animals. Philos. Trans. R. Soc., B, 374:20190088.

78. Lord, J. C., Hartzer, K. L. & Kambhampati, S. 2012. A nuptially transmitted ichthyosporean symbiont of Tenebrio molitor (Coleoptera: Tenebrionidae). J. Eukaryotic Microbiol., 59:246–250.

79. Lotman, K., Pekkarinen, M. & Kasesalu, J. 2000. Morphological observations on the life cycle of Dermocystidium cyprini Cervinka and Lom, 1974, pparasitic in carps (Cyprinus carpio). Acta Protozool., 39:125–134.

80. Mah, J. L., Christensen-Dalsgaard, K. K. & Leys, S. P. 2014. Choanoflagellate and choanocyte collar-flagellar systems and the assumption of homology. Evol. Dev., 16:25–37.

81. Mantzouki, E., Ysnel, F., Carpentier, A. & Pétillon, J. 2012. Accuracy of pitfall traps for monitoring populations of the amphipod Orchestia gammarella (Pallas 1766) in saltmarshes. Estuarine, Coastal Shelf Sci., 113:314–316.

82. Marchant, H. J. & Perrin, R. A. 1990. Seasonal variation in abundance and species composition of choanoflagellates (Acanthoecideae) at Antarctic coastal sites. Polar Biol., 10:499–505.

83. Marques, J. C. & Nogueira, A. 1991. Life cycle, dynamics, and production of Echinogammarus marinus Leach (Amphipoda) in the Mondego estuary (Portugal). Oceanol. Acta, Special issue.

84. Marshall, W. L. & Berbee, M. L. 2010. Population-level analyses indirectly reveal cryptic sex and life history traits of Pseudoperkinsus tapetis (Ichthyosporea, Opisthokonta): a unicellular relative of the animals. Mol. Biol. Evol., 27:2014–2026.

85. Marshall, W. L. & Berbee, M. L. 2011. Facing unknowns: living cultures (Pirum gemmata gen. nov., sp. nov., and Abeoforma whisleri, gen. nov., sp. nov.) from invertebrate digestive tracts represent an undescribed clade within the unicellular Opisthokont lineage Ichthyosporea (Mesomycetozoea). Protist, 162:33–57.

86. Marshall, W. L., Celio, G., McLaughlin, D. J. & Berbee, M. L. 2008. Multiple isolations of a culturable, motile Ichthyosporean (Mesomycetozoa, Opisthokonta), Creolimax fragrantissima n. gen., n. sp., from marine invertebrate digestive tracts. Protist, 159: 415–433.

87. Mendoza, L., Taylor, J. W. & Ajello, L. 2002. The Class Mesomycetozoea: a heterogeneous group of microorganisms at the animal-fungal boundary. Annu. Rev. Microbiol., 56:315–344.

88. Minh, B. Q., Nguyen, M. A. T. & von Haeseler, A. 2013. Ultrafast approximation for phylogenetic bootstrap. Mol. Biol. Evol., 30:1188–1195.

89. Molnár, K. & Székely, C. 2014. Tissue preference of some myxobolids (Myxozoa: Myxosporea) from the musculature of European freshwater fishes. Dis. Aquat. Org., 107:191–198.

90. Morado, J. F. 2011. Protistan diseases of commercially important crabs: a review. J. Invertebr. Pathol., 106:27–53.

91. Morgan, J. A., DeJong, R. J., Jung, Y., Khallaayoune, K., Kock, S., Mkoji, G. M. & Loker, E. S. 2002. A phylogeny of planorbid snails, with implications for the evolution of Schistosoma parasites. Mol. Phylogenet. Evol., 25:477–488.

92. Mylnikov, A. P., Tikhonenkov, D. V., Karpov, S. A. & Wylezich, C. 2019. Microscopical Studies on Ministeria vibrans Tong, 1997 (Filasterea) Highlight the Cytoskeletal Structure of the Common Ancestor of Filasterea, Metazoa and Choanoflagellata. Protist, 170:385–396.

93. Nguyen, L. T., Schmidt, H. A., Von Haeseler, A. & Minh, B. Q. 2015. IQ-TREE: a fast and effective stochastic algorithm for estimating maximum-likelihood phylogenies. Mol. Biol. Evol., 32:268–274.

94. Olson, R. E., Dungan, C. F. & Holt, R. A. 1991. Water-borne transmission of Dermocystidium salmonis in the laboratory. Dis. Aquat. Org., 12:41–48.

95. Owczarzak, A., Stibbs, H. H. & Bayne, C. J. 1980. The destruction of Schistosoma mansoni mother sporocysts in vitro by amoebae isolated from Biomphalaria glabrata: an ultrastructural study. J. Invertebr. Pathol., 35:26–33.

96. Paps, J., Medina-Chacón, L. A., Marshall, W., Suga, H. & Ruiz-Trillo, I. 2013. Molecular phylogeny of unikonts: new insights into the position of apusomonads and ancyromonads and the internal relationships of opisthokonts. Protist, 164:2–12.

97. Paps, J. 2018. What makes an animal? The molecular quest for the origin of the animal kingdom. Integr. Comp. Biol., 58:654–665.

98. Parra-Acero, H., Ros-Rocher, N., Perez-Posada, A., Kożyczkowska, A., Sánchez-Pons, N., Nakata, A., Suga, H., Najle, S. R. & Ruiz-Trillo, I. 2018. Transfection of Capsaspora owczarzaki, a close unicellular relative of animals. Development, 145, e162107. doi:10.1242/dev.162107

99. Patterson, D. J., Nygaard, K., Steinberg, G. & Turley, C. M. 1993. Heterotrophic flagellates and other protists associated with oceanic detritus throughout the water column in the mid North Atlantic. J. Mar. Biol. Assoc. U. K., 73:67–95.

100. Pekkarinen, M., Lom, J., Murphy, C. A., Ragan, M. A. & Dykova, I. 2003. Phylogenetic position and ultrastructure of two Dermocystidium species (Ichthyosporea) from the common perch (Perca fluviatilis). Acta Protozool., 42:287–307.

101. Pekkarinen, M. & Lotman, K. 2003. Occurrence and life cycles of Dermocystidium species (Mesomycetozoa) in the perch (Perca fluviatilis) and ruff (Gymnocephalus cernuus)(Pisces: Perciformes) in Finland and Estonia. J. Nat. Hist., 37:1155–1172.

102. Pereira, C. N., Di Rosa, I., Fagotti, A., Simoncelli, F., Pascolini, R. & Mendoza, L. 2005. The pathogen of frogs Amphibiocystidium ranae is a member of the order Dermocystida in the class Mesomycetozoea. J. Clin. Microbiol., 43:192–198.

103. Raffel, T. R., Bommarito, T., Barry, D. S., Witiak, S. M. & Shackelton, L. A. 2008. Widespread infection of the Eastern red-spotted newt (Notophthalmus viridescens) by a new species of Amphibiocystidium, a genus of fungus-like mesomycetozoan parasites not previously reported in North America. Parasitology, 135:203.

104. Ragan, M. A., Goggin, C. L., Cawthorn, R. J., Cerenius, L., Jamieson, A. V., Plourde, S. M., Rand, T. G., Söderhäll, K. & Gutell, R. R. 1996. A novel clade of protistan parasites near the animal-fungal divergence. Proc. Natl. Acad. Sci. U. S. A., 93:11907–11912.

105. Raghu-Kumar, S. 1987. Occurrence of the thraustochytrid, Corallochytrium limacisporum gen. et sp. nov. in the coral reef lagoons of the Lakshadweep Islands in the Arabian Sea. Bot. Mar., 30:83–90.

106. Rambaut, A. 2017. FigTree v. 1.4.3, a graphical viewer of phylogenetic trees. Software in: http://tree.bio.ed.ac.uk/software/figtree.

107. Reynolds, E. S. 1963. The use of lead citrate at high pH as an electron-opaque stain in electron microscopy. J. Cell Biol., 17:208.

108. Richter, D. J., Fozouni, P., Eisen, M. B. & King, N. 2018. Gene family innovation, conservation and loss on the animal stem lineage. eLife, 7, e34226. doi:10.7554/eLife.34226

109. Ronquist, F., Teslenko, M., Van Der Mark, P., Ayres, D. L., Darling, A., Höhna, S., Larget, B., Liu, L., Suchard, M. & Huelsenbeck, J. P. 2012. MrBayes 3.2: efficient Bayesian phylogenetic inference and model choice across a large model space. Syst. Biol., 61:539–542.

110. Rowley, J. J., Gleason, F. H., Andreou, D., Marshall, W. L., Lilje, O. & Gozlan, R. 2013. Impacts of mesomycetozoean parasites on amphibian and freshwater fish populations. Fungal Biol. Rev., 27:100–111.

111. Ruiz-Trillo, I., Inagaki, Y., Davis, L. A., Sperstad, S., Landfald, B. & Roger, A. J. 2004. Capsaspora owczarzaki is an independent opisthokont lineage. Curr. Biol., 14:946–947.

112. Ruiz-Trillo, I, Lane, C. E., Archibald, J. M. & Roger, A. J. 2006. Insights into the evolutionary origin and genome architecture of the unicellular opisthokonts Capsaspora owczarzaki and Sphaeroforma arctica. J. Eukaryotic Microbiol., 53:379–384.

113. Ruiz-Trillo, I., Roger, A. J., Burger, G., Gray, M. W. & Lang, B. F. 2008. A phylogenomic investigation into the origin of metazoa. Mol. Biol. Evol., 25:664–672.

114. Sambrook, J., Fritsch, E. F. & Maniatis, T. 1989. Molecular cloning: a laboratory manual. 2nd ed. Cold Spring Harbor Laboratory Press, New York, NY. p. 21–152.

115. Scott, J. J., Budsberg, K. J., Suen, G., Wixon, D. L., Balser, T. C. & Currie, C. R. 2010. Microbial community structure of leaf-cutter ant fungus gardens and refuse dumps. PLoS One, 5, e9922. doi:10.1371/journal.pone.0009922

116. Sebé-Pedrós, A., Burkhardt, P., Sánchez-Pons, N., Fairclough, S. R., Lang, B. F., King, N. & Ruiz-Trillo, I. 2013. Insights into the origin of metazoan filopodia and microvilli. Mol. Biol. Evol., 30:2013–2023.

117. Sebé-Pedrós, A., Degnan, B. M. & Ruiz-Trillo, I. 2017. The origin of Metazoa: a unicellular perspective. Nat. Rev. Genet., 18:498.

118. Shalchian-Tabrizi, K., Minge, M. A., Espelund, M., Orr, R., Ruden, T., Jakobsen, K. S. & Cavalier-Smith, T. 2008. Multigene phylogeny of choanozoa and the origin of animals. PLoS One, 3, e2098. doi:10.1371/journal.pone.0002098

119. Shanan, S., Abd, H., Bayoumi, M., Saeed, A. & Sandström, G. 2015. Prevalence of protozoa species in drinking and environmental water sources in Sudan. BioMed Res. Int., 2015, e345619. doi:10.1155/2015/345619

120. Silva, V., Pereira, C. N., Ajello, L. & Mendoza, L. 2005. Molecular evidence for multiple host-specific strains in the genus Rhinosporidium. J. Clin. Microbiol., 43:1865–1868.

121. Simion, P., Philippe, H., Baurain, D., Jager, M., Richter, D. J., Di Franco, A., Roure, B., Satoh, N., Quéinnec, E., Ereskovsky, A., Lapébie, P., Corre, E., Delsuc, F., King, N., Wörheide, G. & Manuel, M. P. 2017. A large and consistent phylogenomic dataset supports sponges as the sister group to all other animals. Curr. Biol., 27:958–967.

122. Snell, E. A., Furlong, R. F. & Holland, P. W. 2001. Hsp70 sequences indicate that choanoflagellates are closely related to animals. Curr. Biol., 11:967–970.

123. Spanggaard, B. & Huss, H. H. 1996. Growth of the fish parasite Ichthyophonus hoferi under food relevant conditions. Int. J. Food Sci. Technol., 31:427–432.

124. Speer, C. A., Dubey, J. P., McAllister, M. M. & Blixt, J. A. 1999. Comparative ultrastructure of tachyzoites, bradyzoites, and tissue cysts of Neospora caninum and Toxoplasma gondii. Int. J. Parasitol., 29:1509–1519.

125. Stamatakis, A. 2014. RAxML version 8: a tool for phylogenetic analysis and post-analysis of large phylogenies. Bioinformatics, 30:1312–1313.

126. Steele, V. J. & Oshel, P. E. 1987. The ultrastructure of an integumental microtrich sensillum in Gammarus setosus (Amphipoda). J. Crustacean Biol., 7:45–59.

127. Steenkamp, E. T., Wright, J. & Baldauf, S. L. 2006. The protistan origins of animals and fungi. Mol. Biol. Evol., 23:93–106.

128. Stentiford, G. D. & Shields, J. D. 2005. A review of the parasitic dinoflagellates Hematodinium species and Hematodinium-like infections in marine crustaceans. Dis. Aquat. Org., 66:47–70.

129. Stentiford, G. D., Bateman, K. S., Feist, S. W., Chambers, E. & Stone, D. M. 2013. Plastic parasites: extreme dimorphism creates a taxonomic conundrum in the phylum Microsporidia. Int. J. Parasitol., 43:339–352.

130. Stentiford, G. D., Neil, D. M. & Atkinson, R. J. A. 2001. Alteration of burrow-related behaviour of the norway lobster, Nephrops norvegicus during infection by the parasitic dinoflagellate Hematodinium. Mar. Freshwater Behav. Physiol., 34:139–156.

131. Stibbs, H. H., Owczarzak, A., Bayne, C. J. & DeWan, P. (1979). Schistosome sporocyst-killing amoebae isolated from Biomphalaria glabrata. J. Invertebr. Pathol., 33:159–170.

132. Suga, H., Chen, Z., De Mendoza, A., Sebé-Pedrós, A., Brown, M. W., Kramer, E., Carr, M., Kerner, P., Vervoort, M., Sánchez-Pons, N., Torruella, G., Derelle, R., Manning, G., Lang, B. F., Russ, C., Haas, B.J., Roger, A. J., Nusbaum, C. & Ruiz-Trillo, I. 2013. The Capsaspora genome reveals a complex unicellular prehistory of animals. Nat. Commun., 4, e2325. doi:10.1038/ncomms3325

133. Taylor, A. C. 1986. Seasonal and diel variations of some physico-chemical parameters of boulder shore habitats. Ophelia, 25:83–95.

134. Taylor, A. C., Field, R. H. & Parslow-Williams, P. J. 1996. The effects of Hematodinium sp. infection on aspects of the respiratory physiology of the Norway lobster, Nephrops norvegicus (L.). J. Exp. Mar. Biol. Ecol., 207:217–228.

135. Tikhonenkov, D. V., Hehenberger, E., Esaulov, A. S., Belyakova, O. I., Mazei, Y. A., Mylnikov, A. P. & Keeling, P. J. 2020. Insights into the origin of metazoan multicellularity from predatory unicellular relatives of animals. BMC Biol., 18:1–24.

136. Tong, S. M. 1997. Heterotrophic flagellates and other protists from Southampton Water, UK. Ophelia, 47:71–131.

137. Torruella, G., de Mendoza, A., Grau-Bove, X., Anto, M., Chaplin, M. A., del Campo, J., Eme, L., Pérez-Cordón, G., Whipps, C. M., Nichols, K. M., Paley, R., Roger, A. J., Sitjà-Bobadilla, A., Donachie, S. & Ruiz-Trillo, I. 2015. Phylogenomics reveals convergent evolution of lifestyles in close relatives of animals and fungi. Curr. Biol., 25:2404–2410.

138. Torruella, G., Derelle, R., Paps, J., Lang, B. F., Roger, A. J., Shalchian-Tabrizi, K. & Ruiz-Trillo, I. 2012. Phylogenetic relationships within the Opisthokonta based on phylogenomic analyses of conserved single-copy protein domains. Mol. Biol. Evol., 29:531–544.

139. Urrutia, A., Bass, D., Ward, G., Ross, S., Bojko, J., Marigomez, I. & Feist, S. W. 2019. Ultrastructure, phylogeny, and histopathology of two novel haplosporidians parasitising amphipods, and importance of crustaceans as hosts. Dis. Aquat. Org., 136:87–103.

140. Van Overdijk, C. D., Grigorovich, I. A., Mabee, T., Ray, W. J., Ciborowski, J. J. & Macisaac, H. J. 2003. Microhabitat selection by the invasive amphipod Echinogammarus ischnus and native Gammarus fasciatus in laboratory experiments and in Lake Erie. Freshwater Biol., 48:567–578.

141. Vilela, R. & Mendoza, L. 2012. The taxonomy and phylogenetics of the human and animal pathogen Rhinosporidium seeberi: a critical review. Rev. Iberoam. Micol., 29:185–199.

142. Wainright, P. O., Hinkle, G., Sogin, M. L. & Stickel, S. K. 1993. Monophyletic origins of the metazoa: an evolutionary link with fungi. Science, 260:340–342.

143. Wilhelm, R. C., Hanson, B. T., Chandra, S. & Madsen, E. 2018. Community dynamics and functional characteristics of naphthalene-degrading populations in contaminated surface sediments and hypoxic/anoxic groundwater. Environ. Microbiol., 20:3543–3559.

144. Zhang, Z., Schwartz, S., Wagner, L. & Miller, W. 2000. A greedy algorithm for aligning DNA sequences. J. Comput. Biol., 7:203–214.

145. Zíková, A., Hampl, V., Paris, Z., Týč, J. & Lukeš, J. 2016. Aerobic mitochondria of parasitic protists: Diverse genomes and complex functions. Mol. Biochem. Parasitol., 209:46–57.

